# Integration of New Information in Memory: New Insights from a Complementary Learning Systems Perspective

**DOI:** 10.1101/2020.01.17.909804

**Authors:** James L. McClelland, Bruce L. McNaughton, Andrew K. Lampinen

## Abstract

According to complementary learning systems theory, integrating new memories into the neocortex of the brain without interfering with what is already known depends on a gradual learning process, interleaving new items with previously learned items. However, empirical studies show that information consistent with prior knowledge can be integrated very quickly. We use artificial neural networks with properties like those we attribute to the neocortex to develop a theoretical understanding of the role of consistency with prior knowledge in putatively neocortex-like learning systems, providing new insights into when integration will be fast or slow and how integration might be made more efficient when the items to be learned are hierarchically structured. The work relies on deep linear networks that capture the qualitative aspects of the learning dynamics of the more complex non-linear networks used in previous work. The time course of learning in these networks can be linked to the hierarchical structure in the training data, captured mathematically as a set of dimensions that correspond to the branches in the hierarchy. In this context, a new item to be learned can be characterized as having aspects that project onto previously known dimensions, and others that require adding a new branch/dimension. The projection onto the known dimensions can be learned rapidly without interleaving, but learning the new dimension requires gradual interleaved learning. When a new item only overlaps with items within one branch of a hierarchy, interleaving can focus on the previously-known items within this branch, resulting in faster integration with less interleaving overall. The discussion considers how the brain might exploit these facts to make learning more efficient and highlights predictions about what aspects of new information might be hard or easy to learn.

## Introduction

A large body of research supports the view that learning in the brain relies on complementary learning systems. One of these systems, based primarily in the hippocampus and related structures, allows the rapid acquisition of new knowledge; the other, based primarily in neocortex, supports the acquisition of structured knowledge that generally builds up over relatively long time scales. The primary evidence for this view comes from the effect of extensive bilateral lesions of the hippocampus and related structures in patient HM and subsequently in other neuropsychological patients and animals. In humans, these lesions profoundly impair the acquisition of arbitrary new memories, while leaving intact previously acquired knowledge and skills, as well as the ability to gradually acquire new skills and often-repeated factual information.

The investigation of computational models based on multi-layer artificial neural networks provided one possible answer to the question, *why* do we have complementary learning systems in the brain? Early after their introduction, research with these models showed that they could gradually acquire structured knowledge through an interleaved learning process, such that experience with each item occurred repeatedly, interleaved with experience with other items. Several models of this kind were presented in the late 1980’s and early 1990’s, demonstrating how the mastery of structured bodies of knowledge of several different types could occur in this way, including knowledge of the mapping from spelling to sound [1, 2], knowledge of the syntactic structure of sentences [3], knowledge of the mapping from sentences to meaning [4], and knowledge of concrete object semantics [5]. Yet these models had a significant deficiency: McCloskey and Cohen [6] showed that an attempt to teach such models new information quickly led to catastrophic interference with the knowledge previously acquired by the network. In McClelland, McNaughton & O’Reilly [7], we drew on earlier ideas of Marr [8] to propose that complementary learning systems exist in the brain to solve this problem: According to our complementary learning systems theory (CLST), multi-layer neural networks, thought to be similar to the neocortex of the brain, need to be paired with a fast-learning, complementary system inspired by the human hippocampus. Features of the hippocampus, according to the theory, provide a specialized type of neural learning system that could learn new things rapidly, allowing new learning to occur without interfering with the knowledge previously acquired in the cortex-like multi-layered neural network. Simulations in [7] showed that if these newly learned items were reactivated from the hippocampus-like system and used as learning experiences for the cortex-like network, interleaved with ongoing exposure to other information already known to the system, the new information would be gradually integrated into the neocortex-like network, without interference.

The recent explosion of research in artificial intelligence using deep neural networks strongly reinforces and supports the ideas laid out above. These networks have achieved tremendous success in building up systems of structured knowledge, providing breakthroughs in machine vision, language processing, and mastery of the most challenging human-invented strategy games, including go and chess. These networks achieve this success through massively interleaved learning, gradually acquiring their abilities over hundreds of millions of experiences in some cases. Catastrophic interference is an important problem for such networks, and developing methods to allow them to acquire new knowledge quickly without interfering with existing knowledge is an open and important research question for artificial intelligence as well as for understanding biological neural systems [9]. Furthermore, many of the solutions that are being explored (e.g., [10]) rely on complementary learning system-like solutions.

Here, we build on recent developments in the neurobiology of learning and in the mathematical analysis of learning in multi-layer neural networks to advance our understanding of the acquisition of new knowledge and of resulting interference with what is already known. We begin by reviewing recent research demonstrating that new knowledge can sometimes be integrated very rapidly into cortex-like systems, with very little interference with prior knowledge, both in biological and artificial learners. We then present new work drawing on a recently developed mathematical theory of learning in deep neural networks to provide a mathematically explicit characterization of some of the observations from previous work, and to address two additional questions about what happens when we learn something new after building up prior knowledge. Specifically, we will address these questions:

- Are all aspects of new learning integrated into cortex-like networks at the same rate?
- Is it possible to avoid replaying everything one already knows when one wants to learn new things?

The answers to these questions should contribute to our understanding of how biological and artificial learning systems work. In addition, as we shall see, the answers to these questions have practical implications, suggesting experience presentation regimes that could make new learning more efficient in both natural and biological learners. Specifically, we will explore when new learning can proceed rapidly, building efficiently on what has already been learned; and when it must proceed slowly to avoid interference. We will see that some but not all aspects of items to be learned might be integrated without interference, and we will identify ways in which interleaved learning might be optimized using *similarity-weighted interleaved learning* to speed the integration of harder-to-integrate aspects of new information into cortex-like networks, while avoiding catastrophic interference.

### New cortical learning can be fast or slow

A very important development in the literature on the neurobiology of learning occurred with the demonstration of rapid integration of new knowledge into neocortical neural networks in the research reported in two major articles by Tse *et al.* [11, 12]. The work clearly demonstrates the key point that such integration depends on the prior establishment of a structured body of knowledge that the authors called a *schema*, building on the classical ideas of Bartlett [13]. A considerable body of recent work in human cognitive neuroscience also investigates the learning of structured bodies of knowledge and the role of schema consistency [14, 15, 16, 17]. Here we focus on the work of Tse *et al.*, returning to issues arising in the human literature in the *General Discussion*.

In these studies, animals were exposed to a flavor-place associative learning task in a specific, previously unfamiliar spatial setting consisting of a 2×3 meter arena with distinctive intra-maze cues within the arena and extra-maze cues outside the arena. On each training day, each animal received one training episode with each of six arbitrary flavor place associations (Figure 1). Within each episode, the animal received a small pellet of food in a start box that could be placed along any of the four walls of the arena, and then was allowed to forage in the arena, where it could find a food well containing three larger pellets of the same flavor. Animals tended to retrieve one pellet at a time and return with it to consume it in the start box before proceeding out into the arena to retrieve the next pellet, so that each episode provided several training trials with the same flavor-place association. Over several weeks, animals gradually acquired the ability to restrict their search preferentially toward the correct place in the environment. Then, animals participated in a single session in which two new flavor-place associations were introduced. While the flavors were completely arbitrary, each of the new assigned places was chosen to be adjacent to the location of one of the food wells associated with a previously-learned flavor (see Figure). Half of the animals received bilateral lesions to the hippocampus within 48 hours of the flavor-place learning trial, while the other half received sham lesions. After a period for recovery from surgery, animals in both groups were tested on both the original and the novel flavor-place associations. Remarkably, both groups not only retained knowledge of the original flavor-place association, but both groups demonstrated learning of the new flavor-place associations; performance was indistinguishable between the two groups, and did not differ from performance with the original six associations. Importantly, the lesioned group could not acquire a second new pair of flavor-place associations rapidly, while the sham lesioned group acquired the second new pair as well as they had learned the first. Thus, an intact medial temporal lobe was required to support rapid new learning, but integration into neocortical networks (as evidenced by retained knowledge after hippocampal removal) occurred within 48 hours. This rapid acquisition of schema-consistent knowledge contrasted with the performance of the same animals in a new arena with different intra- and extra-maze cues, where they were exposed to six new flavor-place associations in a novel spatial arrangement. Here the control animals learned just as gradually as they did in the first environment, while the animals with hippocampal lesions failed to show any progress in learning the new flavor-place associations. Thus, the findings indicate that animals required a preexisting schema for rapid flavor-place association even with an intact hippocampal network.

**Figure 1.**
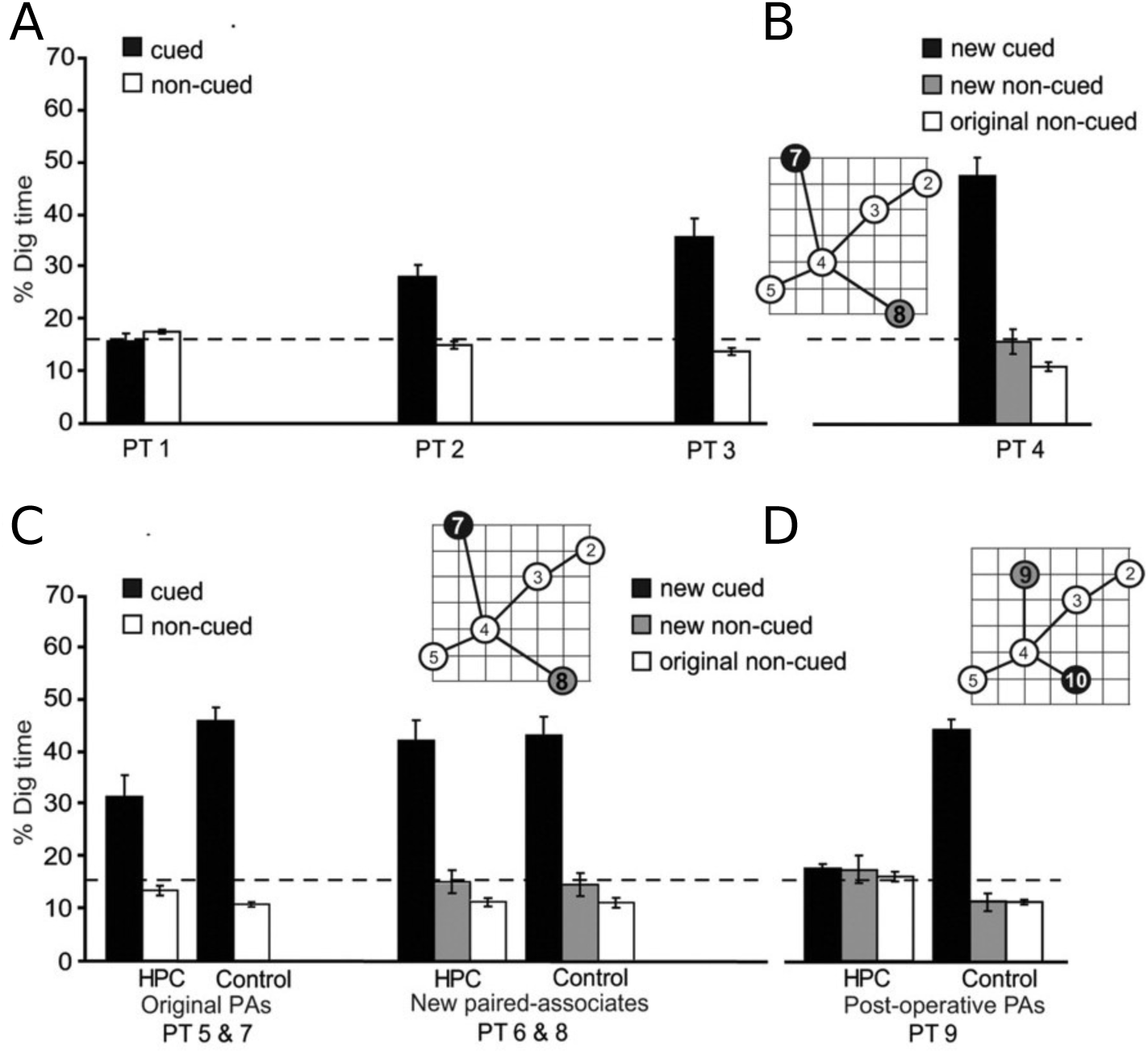
The results of test assessments after different learning experiences as reported in Tse et al. [11]. (A) Shows rats tendency to dig in the correct location when cued with one of the six initially learned flavors associated with the six locations shown in the inset (uncued trials serve as controls) after different numbers of learning sessions (PT1 = 1 session; PT2 = 7 sessions; PT3 = 13 sessions). (B) Shows retention of two new flavor place associations (locations 7 and 8 on inset) one day after one training trial with each association. (C) Shows retention after hippocampal lesions occurring two days after the single learning trial (compared with sham lesioned controls) of the original flavor-place pairs or the new pairs, indicating retention of the new pairs was not hippocampus dependent. (D) Shows complete failure to learn two further pairs (locations 9 and 10) after hippocampal lesion, while controls showed robust learning as expected. From Figure 2, panels B-E, p. 78 of Tse, D., Langston, R. F., Kakeyama, M., Bethus, I., Spooner, P. A., Wood, E. R., Witter, M. P. & Morris, R. G. (2007). Schemas and memory consolidation. *Science, 316* (5821), 76-82. Reprinted with permission from AAAS.

Inspired by these findings and their relevance to CLST, McClelland [18] used the neural network previously used in [7] to show that an analog of the Tse *et al.* findings could be observed in a neural network thought to capture properties of the neo-cortex. We describe this network and several of its characteristics because it provides the basis for much of the new work we describe later in this article.

The network, based on one introduced by Rumelhart & Todd [5], demonstrates how knowledge stored in the connection weights of a multi-layered neural network is gradually acquired through interleaved learning. We had previously used this network in [7] to capture the gradual differentiation of conceptual knowledge that occurs over the course of early through middle childhood as children learn more and more about objects in the world and their properties. Subsequent work [19, 20] showed how this model could account for many aspects of the developmental progression of semantic knowledge acquisition, as well as the disintegration of such knowledge as a function of brain damage, thought to reduce the fidelity of the learned representations of concepts through the random destruction of neurons as a result of neuro-degenerative disease [21].

The network (shown in Figure 2) was initially conceived by Rumelhart & Todd [5] as a way of learning knowledge that might be expressed through propositional statements such as ‘a canary can fly’, ‘a tree is a plant’, ‘a rose is red’ and ‘fish have fins’. The network possessed two pools of input units, the first of which contained a separate input unit for each of eight items (two birds, two fish, two trees, and two flowers), while the second contained a unit for each of four possible semantic relations (ISA, IS, CAN and HAS). The network acquired knowledge of the set of propositions true of each of the eight items by learning from a set of thirty-two training experiences, one for each item-relation pair. When given the item-relation pair as input, the network’s task was to activate the correct corresponding output units, indicating all of the completions that were true for that item and relation. For example, when given ‘robin CAN’ as input, the network’s task was to activate the ‘grow’, ‘move’, and ‘fly’ output units. Training occurred through repeated epochs in which each experience occurred once. After presentation of each item, activation propagated forward through the network, producing activations of units at the representation, integrative hidden, and output or attribute layers of the network. The back propagation learning algorithm [22] was then used to make small adjustments to the connection weights. This is the algorithm still used today in deep learning research to train most deep neural network models in AI and machine learning [23].

**Figure 2.**
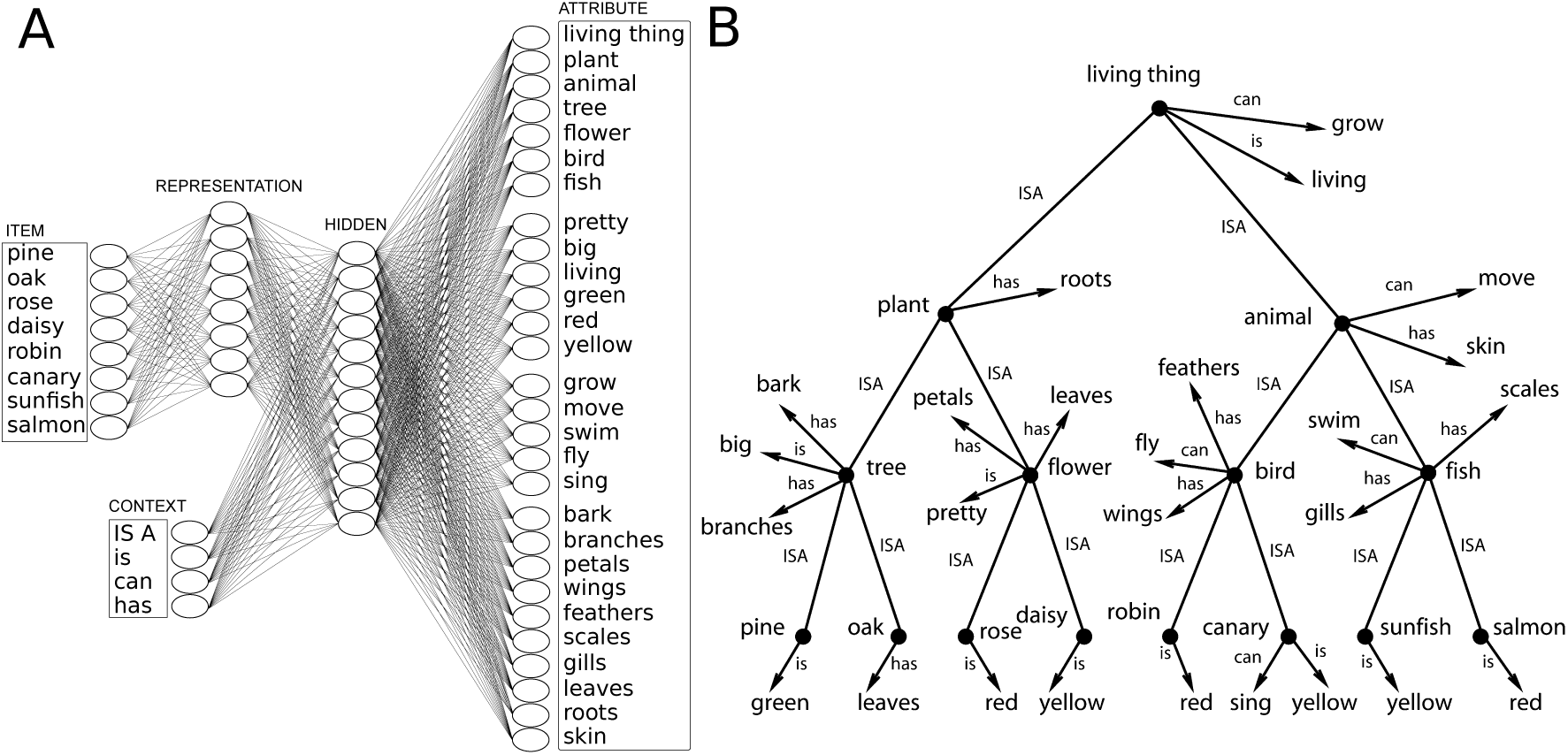
(A) The multi-layer neural network model introduced in [5] and used in simulations reported in [7] and [18]. The full background training set consisted of 32 training items, one for every combination of each of the eight indicated items with each of the four indicated contexts. The network was trained to activate the correct set of output units corresponding to each item-context pair. For example, for *robin can* the network was trained to activate *grow, move, fly*, as indicated in the diagram. (B) An explicit hierarchical tree representing the taxonomic hierarchy or the items used and the set of facts included in the training examples. Facts true of a node higher up the tree (e.g. animal) propagate down to the specific items at the bottom of the tree (e.g. can grow, can move, and can fly propagate down to robin from the living thing, animal, and bird nodes respectively). Adapted with permission from Figure 2, p. 1194 of McClelland, J. L. (2013). Incorporating rapid neocortical learning of new schema-consistent information into complementary learning systems theory. *Journal of Experimental Psychology: General, 142*(4), 1190–1210, American Psychological Association.

As previously demonstrated in [7], the network gradually acquires knowledge of the objects and their properties, in a progressive manner, capturing the progressive differentiation of conceptual knowledge exhibited by children. First the network learns to distinguish the plants from the animals, and at this time in its development, it treats all plants as essentially one kind of thing with one set of properties, and all animals as another kind of thing with another set of properties. Next, it differentiates the birds from the fish, and then the trees from the flowers. Finally, it learns the distinctions among specific items within each of these categories, correctly indicating, for example, that it is only the canary, and not the robin, that can sing. (See [20] for evidence of similar patterns in young children as well as other characteristic patterns exhibited both by developing children and the network).

Once this learning has occurred, the network can be said to have acquired a structured body of knowledge, or in the terminology of Tse et al. [11] a schema. Specifically, following [18], we define a schema as a structure that organizes a body of knowledge so that each item has a well-defined place. The idea is that the rats in the Tse *et al.* experiment learned a spatial schema for the arena in which each of the flavored foods they learned about had a specific place. Likewise the neural network learned a schema – in this case a taxonomic hierarchy – in which each learned item also had a specific place. Following this logic, we can define a schema-consistent item to be an item that could be added to an existing schema without requiring alterations or extensions to it. Examples of schema consistent items include an unfamiliar, but typical bird (labeled a cardinal) and an unfamiliar, but typical fish (labeled a trout). While the real cardinal and trout both have unique differentiating features, the items used in [18] did not; instead, their properties matched those of one of the known birds (a robin) and one of the known fish (the salmon). We see this as corresponding to the situation with the new schema-consistent flavor-place associations in Tse *et al.*, since each of the places used corresponded closely to one of the places used in one of the previously learned flavor-place associations.

To illustrate the importance of consistency with the previously learned schema in the neural network, McClelland [18] also considered an item previously used in [7]. This item, called a penguin, shared the same ISA properties with the known birds, but had the same CAN attributes as the known fish. Crucially, this information is not fully schema consistent, in that it prevents the penguin from being slotted into an existing spot in the learned taxonomic hierarchy.

For each of the new items, McClelland [18] conducted the following simulation experiment. Beginning with the network that gradually acquired the structured knowledge system for the eight initial items, a new input unit was added for the new item. The network was then trained with two training experiences involving the new item, one using the ISA relation and one using the CAN relation. Thus, for the *trout*, the two new items specified *trout ISA LivingThing-Animal-Fish* and *trout CAN Grow-Move-Swim*, and for the *cardinal*, the two new items specified *cardinal ISA LivingThing-Animal-Bird* and *cardinal CAN Grow-Move-Fly*. For the partially schema inconsistent *penguin*, the two new items specified *penguin ISA LivingThing-Animal-Bird* and *penguin CAN Grow-Move-Swim*, crossing the ISA properties of the birds with the CAN properties of the fish. In each of the three simulations, only the two experiences involving one of the new items were presented, without interleaving, making each simulation analogous to the situation in Tse *et al.*, in which learning of new flavor-place associations occurred without interleaving with ongoing exposure to the previously-learned flavor-place associations.

The results were very similar for the case of the cardinal and the trout, so they are averaged together in Figure 3, where they are contrasted with the results for the experiment with the penguin. On the left we show the time-course of learning to activate the correct outputs, as measured by the mean squared error across all of the network’s output units for both item-relation pairs used in the experiment. For both the cardinal and the trout, the error is eliminated very rapidly, indicating that the network very rapidly acquired the ability to learn each of these two items. It should be noted that the network could not have known which set of output units to map the input to – the correct outputs in the case of the trout are not the same as in the case of the cardinal. Thus, in a sense, the network is learning an arbitrary association in each of these two cases, analogous to the arbitrary linking of a specific, previously unfamiliar flavor onto an arbitrary location in the arena. At the same time, however, both the network and the animals in Tse *et al.* [11] are learning something entirely schema consistent, in that both are assigning the new item to an existing place in the previously acquired schema. Indeed, after these learning experiences, the network can infer appropriate fish-like IS and HAS properties for the trout and appropriate bird-like IS and HAS properties for the cardinal when probed for such information.

**Figure 3.**
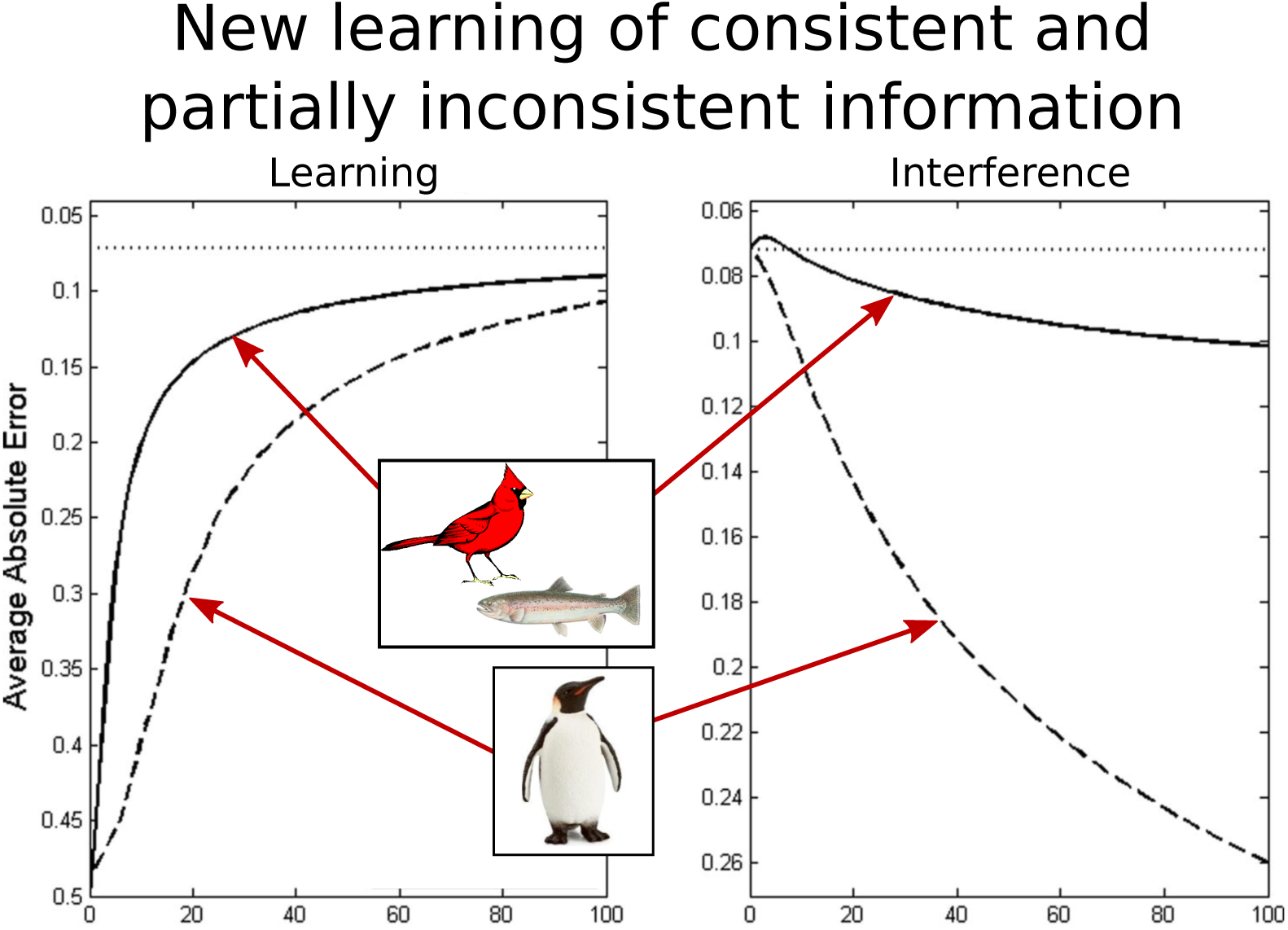
Focused learning of new items either fully or partially consistent with existing knowledge and interference with existing knowledge after learning the eight items and their properties in the network from Figure 2. The left panel shows average network error (reverse plotted; smaller scores are better) for two new items (cardinal and trout) where the new information is fully consistent with existing knowledge and one new item (the penguin) partially inconsistent with existing knowledge. The right panel shows network error on the eight previously learned items as the network learns about the consistent and partially inconsistent items. Fully consistent items are learned far more quickly and with much less interference with prior knowledge. Adapted with permission from Figure 3, p. 1196 of McClelland, J. L. (2013). Incorporating rapid neocortical learning of new schema-consistent information into complementary learning systems theory. *Journal of Experimental Psychology: General, 142*(4), 1190–1210, American Psychological Association.

On the right we show the time-course of interference with other items that results from this rapid learning of the schema-consistent cardinal and trout items. It will be seen that this learning occurs with only the slightest of interference with the existing items. This result is consistent with the finding in Tse *et al.*, that learning the two new flavor-place associations did not interfere with previously acquired flavor-place associations in the same environment.

The very rapid learning of the new schema-consistent items by the network might correspond to learning that occurs in the animals through a combination of direct, within-experience learning and replay-based learning both within the in-arena learning episode and during subsequent off-line waking and resting periods, and the likelihood of this replay may depend on the prior existence of a schema for the environment. Based on the results from the somewhat analogous spatial learning studies of Pfeiffer [24], we know that animals engage in extensive preplay of neural representations of planned trajectories through space as they search for food at previously-learned locations, and that animals also engage in off-line replay during rest and sleep during subsequent offline periods after spatial learning [25]. Of course, from available evidence, the extent of this replay-based learning cannot be known. We can suggest, however, that the failure of animals lacking a hippocampus to acquire new schema-consistent associations within the familiar environment might be a consequence of the failure of the availability of a representation in the hippocampus that would support replay of the correct route back to the target location. We conjecture that these replay events are crucial for allowing the animal to return to the correct location once found, allowing the animal to learn efficiently within the learning episode and allowing for amplification of neocortical learning opportunities through replay both during and after the episode.

Before leaving our consideration of the learning of new schema-consistent items, we can consider how it is that these items can be learned so rapidly, and why it is that this learning produces so little interference. There are basically two points that are essential for understanding these findings. First, the back-propagated error signals arising from the two experiences with each of these items provide strong, coherent, convergent learning signals, driving the input-to-representation weights in Figure 2 coherently toward values that treat the item either as a bird-like animal (in the case of the cardinal) or as a fish-like animal (in the case of the trout). Since (as we will discuss in the theory section of this article below) the learning signals depend on the knowledge already built up in these weights during schema acquisition, they are far stronger after schema acquisition than they would be earlier in learning. This allows the network to exhibit a readiness to learn that depends crucially on the prior acquisition of the relevant schema. Second, because the changes build up so quickly, the error at the output layer is very quickly eliminated, so that there is very little need or opportunity to adjust the connections that are shared among the different concepts in the weights forward of the representation layer in the network.

We now turn attention to the experiment with the penguin. The penguin was the example used in the simulation of interference with previous learning to motivate the Complementary Learning Systems theory in [7]. There, when the network was trained only with the ISA and CAN experiences involving the penguin, this resulted in substantial interference with previously learned items – particularly, the birds and fish. Indeed, we replicate this finding here, as shown in Figure 3. Looking first on the left of the figure, we see that learning proceeds substantially more slowly for the penguin than it does for the trout and the cardinal, and correspondingly, on the right, the penguin produces far more interference with the previously-learned birds and fish. The reasons for these effects can be understood in terms of the ideas we already considered to explain the faster learning and lack of interference for the schema consistent trout and cardinal items. First, the back-propagated error information now partially cancels out, driving the connection weights in one case toward weights that would work for a bird and in the other toward weights that would work for a fish. The result is slower acquisition of connection weights from the input to the hidden layer that assign the penguin a representation intermediate between a fish and a bird. Second, the connection weights forward from the representation layer to the output for this compromise representation will partially activate bird and fish in the ISA context and swim and fly in the CAN context. Adjusting these weights to correctly activate bird for the ISA context and swim for the CAN context will result in a tendency to call all the animals birds (interfering with the known fish) and to say they all can swim and not fly (interfering with the known birds).

It is useful to note that the results we observe with the penguin are not as extreme as the results we might observe if we trained the network with a pair of examples that are completely inconsistent with the learned schema based on the original eight items. Consider the case in which the input X-ISA is paired with three units randomly selected across all the output units of the network, and similarly for the input X-CAN. This would radically disrupt the schema acquired by the network, in which all ISA outputs come from one subset and all CAN outputs come from another, while also cross-pairing features that may never have occurred together in any items. Learning of such items would be very slow, and, as in the simulations reported in early work by McCloskey and Cohen [6], this learning would drastically interfere with the networks knowledge of all items previously acquired (for details, see [18]).

The work reviewed above reflects the complexity of the issues that arise when we consider new learning in deep neural networks – issues that may have analogs in new learning in biological learning systems such as humans and non-human animals. It would be an oversimplification to characterize new learning in a cortex-like neural network as inherently fast or slow – instead, as discussed in [18] such learning has to be seen as prior knowledge dependent. Furthermore, it would be wrong to state that rapid learning in a cortex-like network always produces interference, or that the interference is necessarily completely catastrophic. Such effects are matters of degree, and depend on the consistency of the new information to be acquired with pre-existing knowledge.

### Toward a Theory of Learning in Deep Neural Networks

The analysis we have provided thus far is largely qualitative and intuitive. It has been helpful in clarifying when learning can be fast and slow and when learning can result in interference. However, many questions remain, and the qualitative principles we have described may not provide a sufficient guide to allow predictions about when we will observe rapid learning and/or interference with previously acquired knowledge. In this section, build on the theory presented by Saxe, McClelland & Ganguli [26] to provide a mathematically explicit characterization of some of the ideas we have been discussing, and to address the two additional questions raised in the introduction: *Are all aspects of new learning integrated into cortex-like networks at the same rate? Is it possible to avoid replaying everything one already knows when one wants to learn new things?*

We begin by presenting the basic theory, applying it to understand the dynamics of learning in simplified versions of deep neural networks, including initial learning of a domain of information, and learning something new once initial domain learning is complete. We then return to our two questions after presenting the basic features of the theory.

### Statistical Structure and Dynamics of Learning

In the theory, we rely on a beautiful relationship between a statistical characterization of the structure of the knowledge in a set of experiences and the dynamics of learning this structure in a deep, linear neural network. We will first describe how the theory allows us to characterize the time course of learning from *tabula rasa* – a completely uninformed initial state – as it was developed in [26] Then, we can proceed to apply it to understand the learning of new items, once some prior knowledge has already been acquired. The theory relies on mathematical ideas that may be unfamiliar to many readers. To make it as accessible as possible, we adopt a tutorial approach, in hopes of enhancing engagement between theoretical and experimental research in the neurobiology of learning and memory.

We begin by characterizing the statistical structure in a body of experiences and then proceed to discuss the dynamics of learning this structure in a deep linear neural network.

#### Statistical structure in a set of learning experiences

In characterizing the statistical structure in a set of learning experiences, we focus on data sets that are hierarchically structured as in the data set of living things we have already been considering, although the mathematical theory applies more generally, and can be used to capture a wide range of different types of structure [26]. Specifically, we will focus on sets of items that can be generated by a process that starts at a root node, and then successively branches. Features that appear in the items can be introduced at any node and can occur within any of the node’s children but cannot occur in children of nodes in other branches. The domain of real living things is generally thought of as hierarchically structured in this way, but not as strictly as in data set we focus on here, since sometimes dimensions of variation (for example, gender) can cut across branches of a hierarchical tree. We discuss deviations from the strict hierarchical structure explored here in the general discussion.

For the purpose of our analysis, it is useful to display the items in our data set in the form of an item-property matrix, in which each item occupies a row in the matrix and each property corresponds to a column (Figure 4a). If an item has a property there is a 1 in the corresponding cell of the matrix (shown as black in the figure); otherwise the cell contains a 0 (shown as white).^1^

**Figure 4.**
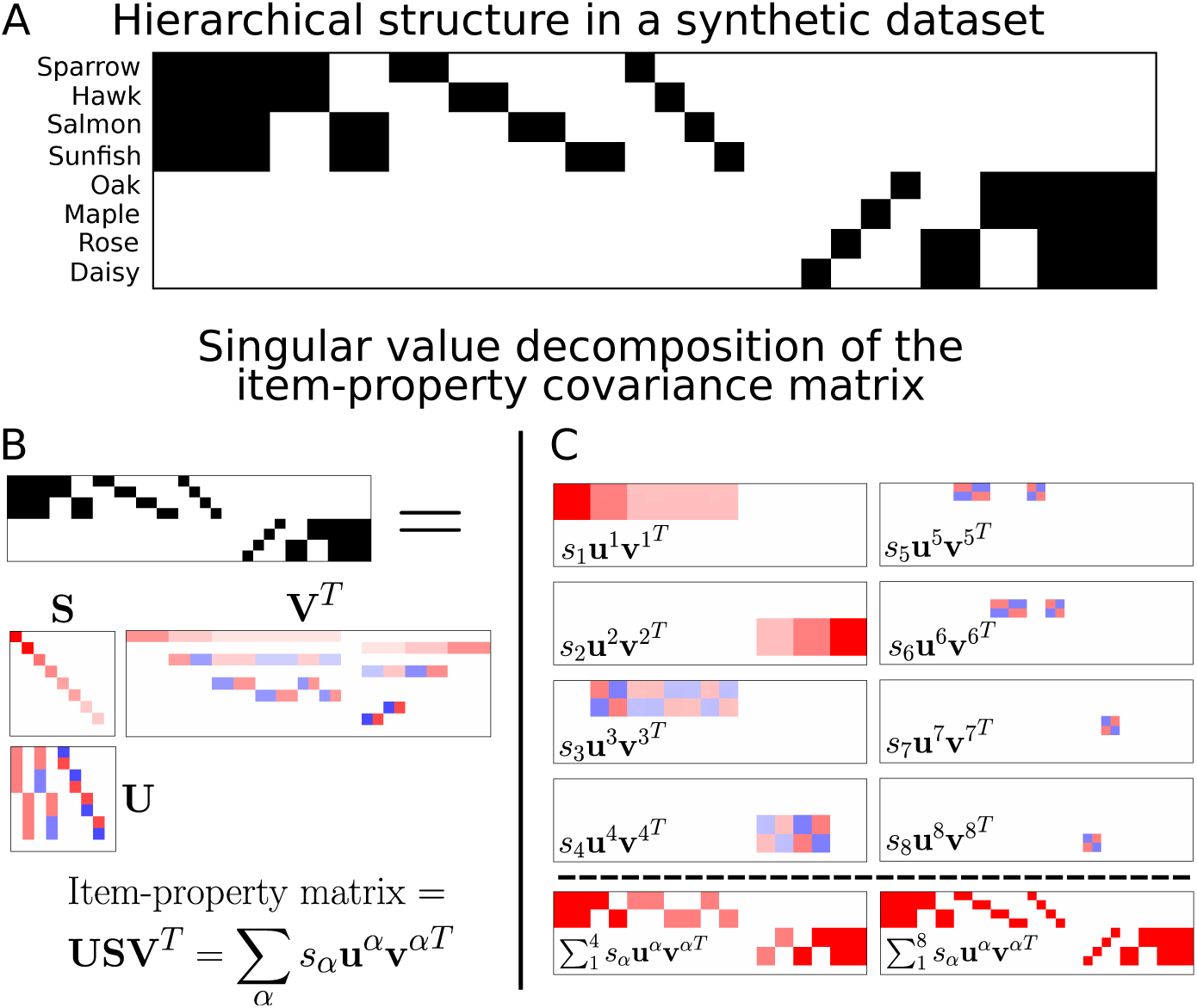
(A) The item-property matrix associated with the new data set used in our new simulations. Each row corresponds to an item, and each column corresponds to a feature (e.g. has eyes or can fly); black indicates that the item has the feature, white indicates that it does not. (B) The item-property matrix is shown again without labels, indicating how it can be decomposed into a set of singular dimensions, each with its own separate strength. Red is used for positive numbers, and blue for negative, with darker shades corresponding to larger values. (C) Each component of the sum in the equation at the bottom of (B) is displayed separately above the dashed line (first 4 components on the left, last 4 on the right). The sum of the first four components is shown below the dashed line on the left, and the sum of all eight, which corresponds to the item-property matrix, is shown below the last line on the right. **Note:** In the matrices **U** and **V**^**T**^ in panel B and in all matrices in panel C, the darkest red corresponds to 1, the darkest blue corresponds to −1, and white corresponds to 0. In the **S** matrix, the darkest red corresponds to 5, and white corresponds to 0.

What one should see in this matrix is that there are two sets of four items, each consisting of features with no overlap at all with the features of the other set of four items. Within each set of four items, there are two sets of two items that share some features but differ on others, and within each pair of items, there are further differentiating features. Although the statistical structure in the matrix can be characterized in completely abstract terms, we support intuition by considering the items to be two birds, two fish, two trees, and two flowers, and consider the properties to be attributes like those we have been considering for such items above. The data set is simpler than the one we have used up to now: we now treat attributes of different types like *can fly* and *has wings* homogeneously, so that each item maps onto a single vector of features.

We have arranged the properties of the items for ease of perusal, placing the features shared by all of the animals on the far left and the features shared by all of the plants on the far right. Proceeding leftward from the shared animal features (which could be attributes like like *can move*, and *has eyes*), we next have features that are present for birds and not for fish (such as *can fly* and *has wings*), then the reverse. Next we see pairs of features that apply to the sparrow only, the hawk only, and each of the two fish only. For example, we think of the hawk as *large and fierce* and the sparrow as *small and meek*. We then encounter four more features, each of which only occurs in one item. We might think of this as a feature that might differentiate it from all other items or that might correspond linguistically to the item’s name. Further to the right, we see a set of unique identifying features, one for each plant, then features that apply only to flowers, features that apply only to trees, and finally the previously mentioned features that apply to all of the plants.

To characterize the structure in the item property matrix, we rely on the *singular value decomposition* (SVD), an analytic method related to Principal Components Analysis (PCA).^2^ The SVD decomposes the item-property matrix into a set of dimensions indexed by *i*. Each dimension is characterized by an item classifying vector **u**^*i*^, a feature-synthesizing vector **v**^*iT*^, and a positive scale factor called the singular value *s*_*i*_.^3^ In the figure, the **u**^*i*^ appear as the columns of the matrix **U**; the **v**^*iT*^ appear as the rows of the matrix **V**^*T*^, and the *s*_*i*_ appear as the diagonal entries in the matrix **S** as shown for our data set in Figure 4B. Rather than display numbers in these matrices, we use red for positive values, blue for negative values and white for 0. Darker red or blue color corresponds to greater magnitude. For the singular values, the darkest red corresponds to the value 5; otherwise, the darkest red and blue correspond to 1 and −1 respectively.

The dimensions are all mutually orthogonal, and the **u**^*i*^ and **v**^*i*^ all have unit length, allowing each *s*_*i*_ to correspond to the overall strength of the corresponding dimension. The first dimension is defined as the dimension that captures the largest possible amount of the total variance (sum of the squares of the values in all of the cells) in the item-property matrix in another matrix that can be constructed by taking the outer product of two vectors – this is the best we can do with just one item classifying vector and one feature synthesizing vector. Each successive dimension is then the one that captures the largest possible amount of the remaining variance after all of the previous dimensions has been removed (there can be ties, as here, and in that case the choice of which to remove first is arbitrary). In this context, a useful aspect of the singular value decomposition is that the scale factor *s*_*i*_ corresponds to the square root of the variance in the data that the dimension explains.

It is important to see that the singular value decomposition captures the hierarchical structure in the data. That is, each dimension is a matrix that captures information about items within a particular branch of the hierarchy. The matrix for each dimension is formed by taking the outer product of its item-classifying vector **u**^*i*^ and its feature-synthesizing vector **v**^*iT*^, and scaling this by the singular value *s*_*i*_, written as *s*_1_**u**^1^**v**^1*T*^. We have shown the matrix for each of the eight dimensions in our data set in Figure 4C. We can see that the first dimension as telling us the average values of the features of the animals; they all share the first four features so the average value for these features is 1; two of the four have each of the next four features so the average value for these is .5; and one of the four has each of the remaining features, so the average value for these is .25.

Thus far, we have seen that designating the first half of the items as members of the same class and assigning all of them the average of their feature values captures more of the variance in our data set than we could capture with any other choice of one item-classifying vector and one feature-synthesizing vector. If we look at the matrix for the next dimension, *s*_2_**u**^2^**v**^2*T*^, we see that designating the second half of the items as members of a different class and assigning all of them the average of their feature values is the best we can do to capture the remaining variance in our data set with one item classifying vector and one feature synthesizing vector. Taking the first two dimensions together, we see that just knowing how to split the items into two sets, and knowing the average feature values of the items in each of the two sets, already captures quite a lot of what there is to capture about our data set.

The remaining dimensions serve to capture the successive branching structure of our hierarchy. The third dimension captures the split among the animals into two birds and two fish. This dimension, *s*_3_**u**^3^**v**^3*T*^, has positive and negative values that produce offsets from the mean values averaged over all four animals as reflected in the first dimension. To see how this works, note that the third item classifying vector **u**^3^ is positive (red) for the birds but negative (blue) for the fish, while the feature synthesizing vector **v**^3*T*^ is positive (red) for the properties true of birds and not fish, and negative (blue) for properties true of fish but not birds. The outer product of these two vectors, when scaled by the third singular value, produces the matrix *s*_3_**u**^3^**v**^3*T*^, and when this matrix is added to the *s*_1_**u**^1^**v**^1*T*^ matrix, the result is a matrix containing the average features of the birds in its top two rows and the average features of the two fish in the next two rows. The fourth dimension captures the corresponding offsets from the matrix of average plant values for the two flowers and the two trees in a similar way. We show the sum of the first four dimensions in Figure 4b below the individual matrices for dimensions 1 to 4. Once again, this is the best we can do in capturing the variance in the data with four dimensions each consisting of one item-classifying vector and one feature synthesizing vector. With these four dimensions, we can classify the items into four groups of two items, capturing the average properties of the items in each of these four groups.

We are still missing out on the ways in which the individual items differ from each other within each of the four mid-level categories. For that, we need four additional dimensions. The fifth and sixth equal strength dimensions capture the offsets to the two average bird vectors needed to fully capture the features of the sparrow and the hawk and the offsets to the two average fish vectors to capture the salmon and sunfish. The remaining dimensions play the same roles for the four plants.

We display the sum of the eight matrices below the individual matrices for dimensions 5 to 8, and we see that we have exactly reconstructed our original item-property matrix. In general, for hierarchically structured data sets, where the same number of new features is added with each categorical split, the successive dimensions will capture average values starting from the highest level, and one dimension is required to capture each pairwise categorical split (for an *N*-way split at the same categorical level, *N* − 1 new dimensions are required). See [26], for details and boundary conditions. In our case, the dimensions on the animal side are stronger because more features (each adding variance) lie on the animal side.

#### Dynamics of learning from a random initial state

Thus far, we have simply characterized the structure in our item-property matrix, but have not yet considered how this structure is learned in a neural network. We next consider what happens when this structure is acquired in the deep, linear neural network shown in Figure 5. This network can be seen as a simplified version of the Rumelhart network shown previously in Figure 2, with the relation units removed, so that the network contains a single input unit for each item and a single output unit for each feature. We call this network *linear* because the output of each unit at a given layer is just a linear function of the activations of the units at the preceding layer (that is, it is simply the sum across all of the units at the preceding layer of the product of the activation of the unit times the corresponding connection weight). We call the network *deep*, because it includes a layer of hidden units between the input and the output layer, and because the characteristics we describe are also exhibited when there are more hidden layers. Although the computations that can be performed by a deep linear network can also be performed by an equivalent network that directly links input units to output units with a single layer of connection weights, the dynamics of learning in deep linear networks are surprisingly non-linear and very different from those of a network without a hidden layer [26]. The key insight here is that the signals required to drive learning in the weights to the hidden layer from the input layer depend on the existing knowledge in the weights to the output layer from the hidden layer, and vice-versa. For mathematical precision, these relationships are captured in the vector-matrix equations shown in Figure 5. Here we convey the crux of the idea while avoiding heavy reliance on the conventions of the linear algebra. First, note that, according to the back-propagation learning rule, the change to each weight is given by the learning rate *λ* times the activation on the input side of the weight, times the error signal on the output side of the weight. For the hidden to output weights **W**^2^, the error signal is the vector of differences **y**′ between the network’s output **ŷ** and the target vector **y**, and the activation signal on the input side is the hidden layer activation pattern **h**, which depends on the input to hidden weights **W**^1^. If these input to hidden weights were all zeros, the hidden layer activations would all be 0, so there could be no change to the hidden-to-output weights. For the input to hidden weights, the activation on the input side is just the input pattern *x*, and the error signal on the output side of these weights is the learning signal *y*′ back-propagated through (i.e. mutiplied by) the hidden-to-output weights. So, if these hidden-to-output weights were all 0, there would be no change to the input-to-hidden weights.

**Figure 5.**
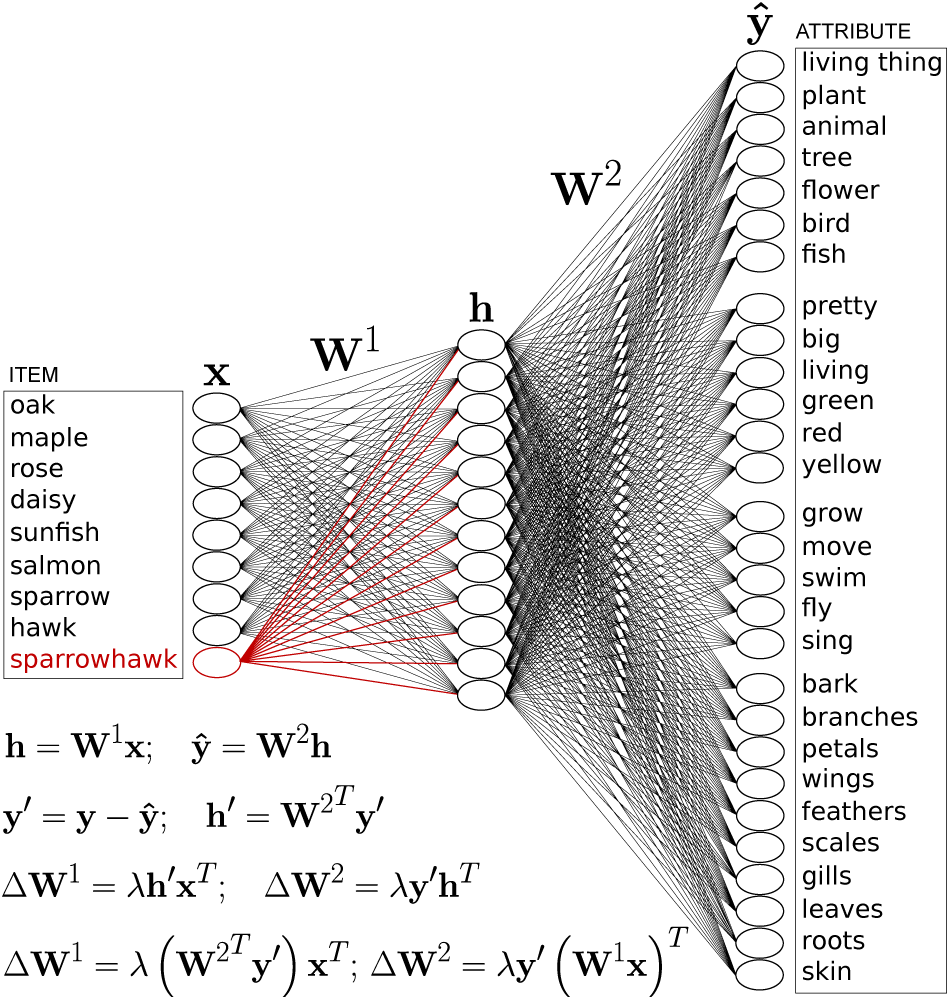
The simplified network architecture used in our analysis and simulations, showing an input unit for each of the eight base items in the new data set, plus an additional, initially unused input unit for sparrowhawk, to be learned after learning the eight base items. There is now only one hidden layer, and the network is linear, in that the vector of activations of hidden units of the network **h** is just the product of the input vector times the input-to-hidden matrix **W**^1^, and the network output **ŷ**, is just the product of the vector **h** times the hidden-to-output weights **W**^2^, as shown in the first row of inset equations. When the network is trained using back-propagation, the error signal **y**′ for the hidden-to-output weights is the difference between the target vector **y** and the network’s output **ŷ**, while the error signal **h**′ for the input to hidden weights is the output-layer error signal **y**′ back-propagated through the hidden to output weights **W**^2^, as shown by the second row of equations. The third row of equations expresses the fact that the change in each weight matrix depends on the learning rate *λ* times the activations at the input to the matrix times the error signal at the output of the matrix. The fourth row of equations substitutes factors from the first and second rows to make explicit how the change to each weight matrix depends on the other weight matrix.

More generally, the learning in each weight matrix depends on the knowledge already stored in the other matrix. If the weights in either matrix are very small, learning in the other matrix will proceed slowly. As the weights in each matrix begin to build up, learning will proceed more quickly in the other, resulting in an acceleration of learning that eventually slows down as all of the error in the mapping from input to output is eliminated. When there are more than two layers of weights, learning in each layer depends on *all* of the other layers of weights. A further important point is that the dynamics of learning we shall be exploring in our simplified network are qualitatively similar to the dynamics of learning in the highly non-linear deep network we first explored in [7] and reviewed above. Indeed, the analysis we describe here was developed in [26] to provide a theoretical understanding of patterns that had previously only been observed in simulations.

The truly remarkable fact about the dynamics of learning in a deep linear network is that it is completely characterized by the SVD, subject to an influence of the initial values of the connection weights at the beginning of the learning process. That is, if the training of the network proceeds as in our earlier simulations with the original Rumelhart network, such that each item is presented once in each training epoch, the network’s input-output (*IO*) matrix at a particular time *t* measured in epochs will be characterized by the two equations below and the corresponding curves presented in the top panel of Figure 6.

**Figure 6.**
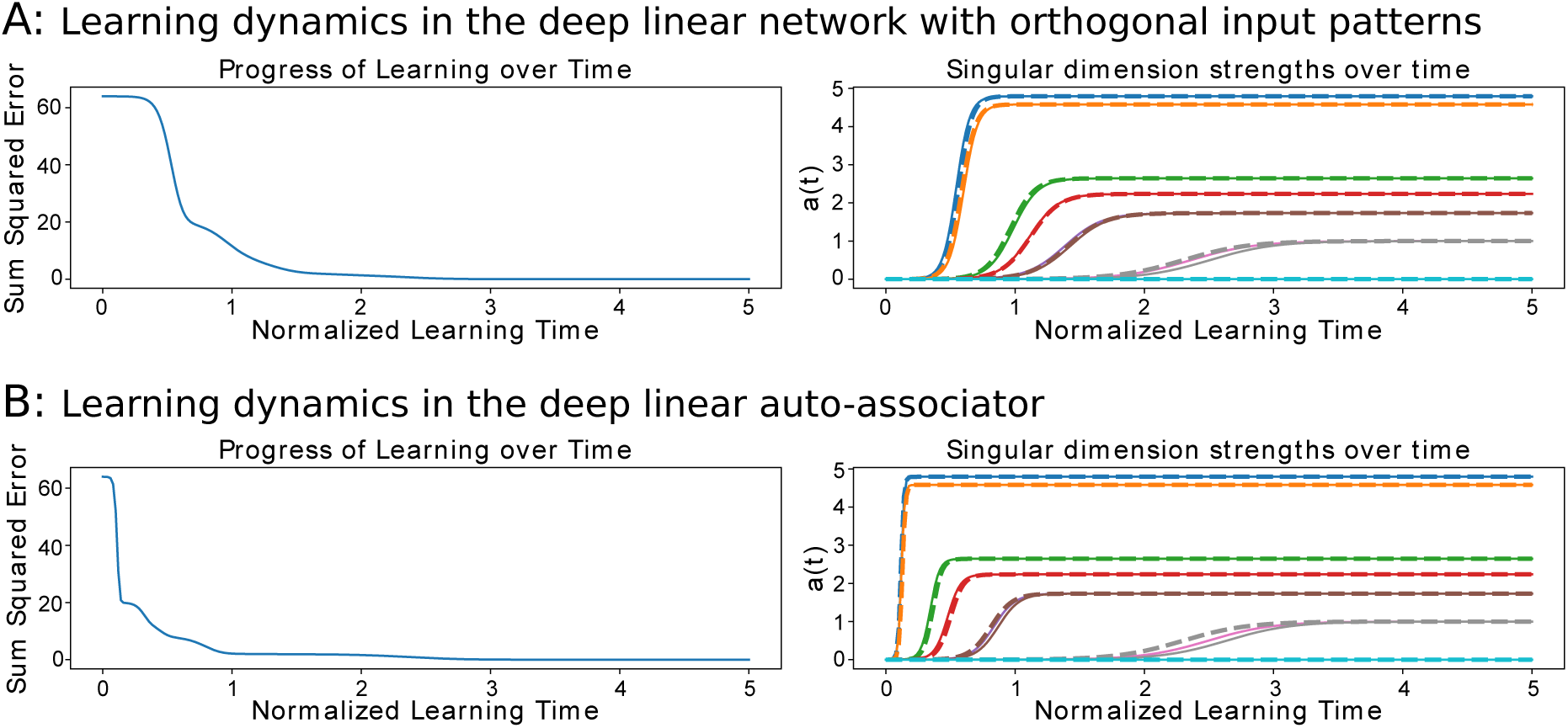
A: The time course of learning in the deep neural network with orthogonal (one-hot) input patterns, as shown in Figure 5, trained on the eight-item data set shown in Figure 4. B: The time course of training on the same items in a deep auto-associative network, where the input pattern is the same as the target output pattern. On the left, the sum squared error over all 8 patterns is shown as a function of normalized training time. On the right, the singular value dimension strengths calculated from the network’s output are shown using solid lines; the mathematically expected curves are shown as dashed lines. Discrepancies between observed and expected vary from run to run of the simulation and are due to variation in the projections of the random initial weights in the network onto the dimensions of the SVD of the item-property matrix. In the one-hot case, time to learn a dimension is inversely proportional to the strength of the dimension in the training set. In the auto-associator case, time to learn is inversely proportional to the square of the strength of the dimension.

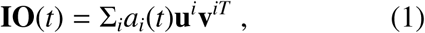

where

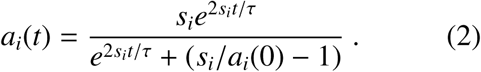

Let us understand what these equations mean. The matrix **IO**(*t*) is the set of output vectors produced by the network where each row vector corresponds to the output produced for one of the inputs to the network. If each *a*_*i*_(*t*) value was equal to *s*_*i*_, then the network’s output would correspond to the item-property matrix. What the first equation expresses, then, is that the output of the network at time *t* can be described as the weighted sum of the dimensions of the SVD of the training data. What the second equation captures is the fact that the weight on each dimension follows a sigmoid curve starting from an initial value *a*_*i*_(0) to its asymptotic value *s*_*i*_.

Figure 6 shows the values computed from the theory for these equations as a function of *t* for the choice *s*_*i*_(0) = .001 and for a value of *τ* dependent on the network learning rate, as described in Supplementary Materials, Section I. We represent *t* in normalized time units corresponding to the number of epochs, or complete sweeps through the training set, times the learning rate, which we choose very small to produce a faithful approximation to the continuous learning equations. This allows us to capture the continuous nature of the learning and to highlight that the choice of the actual learning rate simply determines the time scale of the process. These curves are superimposed on a plot of the actual dynamics of learning in a simulation of the neural network shown in Figure 7. For the simulation, we employ a network with 32 hidden units, initializing the weights with small random values such that their SVD can be characterized by 32 random initial dimensions with an average initial strength of .001. The observed values of the *a*_*i*_(*t*) are then determined by actually applying the SVD to the network’s output at each time *t* and plotting the corresponding simulated values in the figure. The small deviations between network model and theory are due to effects of the random initial values of the connection weights, which affect the true values of the *a*_*i*_(0), which are approximated by fixed constant values in the equation.^4^

**Figure 7.**
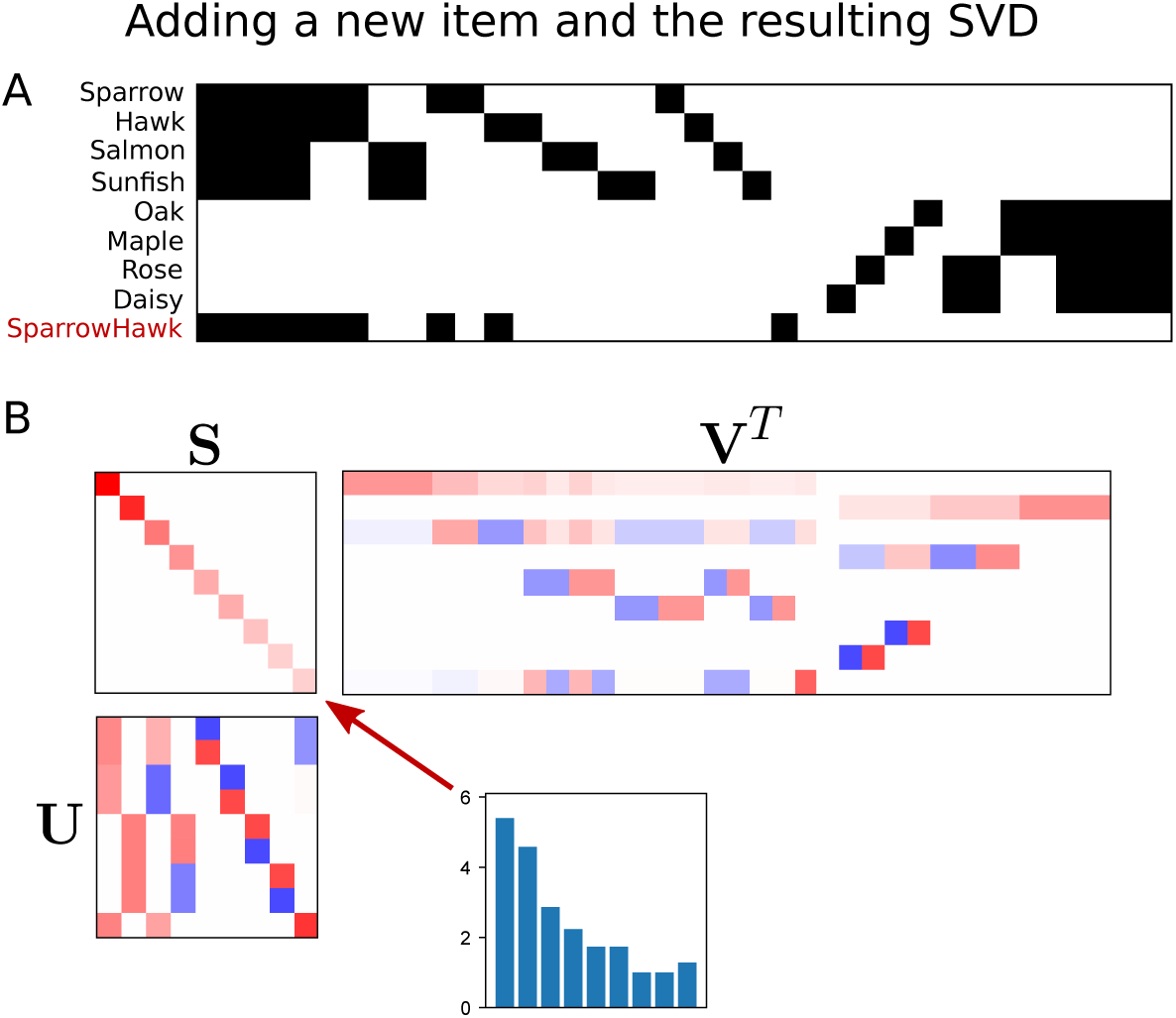
(A) The augmented data set containing the sparrowhawk in addition to the existing eight items from the training set presented in Figure 4. The sparrowhawk is small like the sparrow but fierce like the hawk, and also has its own unique identifying feature like all of the other items. (B) The SVD of the new full data set. Comparing this with the SVD of the original data set, dimensions 1 and 3 have been slightly altered, as can be seen by looking at the top and 3rd rows of matrix **V**^*T*^, and are slightly stronger, since there is another item whose statistics are contributing to the representation of all of the animals (dimension 1) and to the representation that differentiates the birds and fish (dimension 3). In addition there is a new added dimension (last row of **V**^*T*^), required to allow the network to correctly map all three birds to their appropriate output features (dimension 5 still distinguishes between the sparrow and the hawk as it did previously). We have placed this dimension last in the SVD to make it easy to facilitate comparisons with the eight item case and to see how the sparrowhawk projects onto these dimensions (shown in the bottom row of the **U** matrix). (Apparent changes to dimensions 7 and 8 are meaningless rotations; their outer products *s***uv**^*T*^ have not changed.)

For our purposes, the most important fact to come out of this analysis is the observation that the learning proceeds according to the hierarchical structure of the training data, as captured by the SVD. The time course of learning of each dimension of the SVD follows a sigmoidal curve whose parameters depend only on the overall strength of the dimension *s*_*i*_ and its initial value *a*_*i*_(0). Three specific points are relevant here. First, the time required to learn about a particular hierarchical split in the data (as characterized, say, by the time required to reach 1*/*2 of the corresponding dimension’s final value), depends primarily on the strength of the dimension, or the amount of variance in the data the dimension accounts for. Second, the sigmoidal character of the learning of each dimension can have the consequence of producing a highly stage-like process, so that the network can have mastered some dimensions fully (in our case, for example, the distinctions between plants and animals, captured in the strongest two dimensions in the SVD) before it exhibits any appreciable sensitivity to details of the differences among particular items (captured in the weakest dimensions). Taking these two points together, we see that detailed information about particular items captured by weaker dimensions in the training data is learned much more slowly than general information that is shared across many items. This pattern is also exhibited in human development and, as previously mentioned, in the dynamics of the deep nonlinear networks we first explored in [7]. These networks progressively differentiate hierarchies like the hierarchy of plants and animals, starting with the highest level categorical splits (for results comparable to those in Figure 6A, see Figure 4C of [19] or Figure 3.4 of [20]. Our analysis of deep linear networks thus seems relevant to understanding the dynamics of learning in more complex real and artificial neural networks.

The final point is that the time course of learning about each dimension depends on what is already known about this dimension at the time we begin to observe the process of learning, as reflected in the quantity *a*_*i*_(0). That is, *the time required to learn about each dimension depends on what we already know about it at the time we start to measure learning*. This observation is particularly relevant to our purpose in the next section, which is to characterize the time course of new learning from any given point we might choose to define as the reference time *t* = 0, as we shall see after a brief consideration of extensions of the domain of the theory.

#### Extending the theory to experiences with other forms of inputs

A feature of the simulations we have considered thus far is that they rely on a ‘one-hot’ input representation of each item such that each item is represented by a single neuron. If this were a real limitation of these networks it would be deeply disappointing, since it seems unlikely that biological systems would actually rely on single, grandmother-cell neurons. Happily, it turns out that this is not a real limitation. Here we briefly consider three alternative cases. The first is the case in which the input vectors are not onehot vectors, but instead are multi-dimensional orthogonal vectors, each of equal magnitude (where the magnitude is the square root of the sum of the squares of the values in the vector). The theory is completely unchanged in this case, although the actual input-classifying vectors will not be as transparently identifiable with the conceptual identity of each of the items. Indeed the one-hot version of the theory can simply be seen as a convenient transformation of the input patterns into a basis that allows the item classification vectors to be rendered more interpretable than they would be if they remained in the actual input coding space. Thus, for example, if the neural representations of the odor stimuli used in the Tse *et al.* experiments were approximately orthogonal high-dimensional vectors, then this input representation would be a useful way of modeling how they function as input to a deep neural network that maps them onto a structured system of internal representations of corresponding places in an arena.

The second case we consider is one in which the patterns correspond to vectors that may have some degree of correlation, or similarity, to each other. Such correlations will induce a tendency for what is learned about one input to transfer to similar items, and this is likely to be helpful if, as is often the case, items that appear similar share other characteristics, or if the very same item appears slightly differently to the senses on different occasions. The situation can be more problematic if similar items must be mapped to completely different outputs, as could be the case if, for example, two very similar flavors in the Tse *et al.* experiment had to be mapped to two distinct places in the environment. The theory can address the effects of such correlations (and can capture differences in strength of input activations, capturing aspects of differences among features in their perceptual salience) as long as the set of patterns to be mapped to distinct outputs are linearly separable. (In cases where the inputs themselves are not linearly separable, the brain may employ conjunctive encoding schemes to help ensure that the patterns that are input to cortical learning systems are linearly separable; however, the theory does not address the dynamics of learning to form such a linearly separable code).

Third, we consider the case in which the neural network is a deep linear auto-associator, i.e., a linear neural network that simply associates each pattern with itself, via a layer of hidden units (rather than relying on direct unit-to-unit connections). The auto-associative case is interesting because this form of learning is entirely driven by the distribution of experiences, without requiring separate input and target patterns to be provided by the environment, and indeed without requiring the experiences to be labeled as falling into distinct categories in any way. Instead of having one-hot input units for each item, the patterns corresponding to the items in the training set serve both as inputs to and targets for learning in the network, and the task of the network is to learn an input connection weight matrix W1 that maps from the input to the hidden layer and an output connection weight matrix W2 that maps from the hidden layer to reproduce the input on the output. Non-linear networks of this type have been used extensively in deep learning research, where the hidden layer representations are thought of as providing a compressed, invertible representation of the input [28]. The auto-associator is also interesting because it can perform pattern completion, a general form of memory in which any fragment or approximation of an input pattern can be thought of as serving as a potential cue for the reconstruction of all aspects of the pattern. Our linear auto-associator is also interesting in that it can be considered to capture one step of a recurrent computation in which the output of the auto-associator is repeatedly fed back into itself, allowing the learned states in the network to function as attractors.

When we train such a network with our data set, the time course of the development of the *IO* matrix (as before, the matrix of outputs produced by each of the eight input items) is characterized by the curves shown in the bottom panel of Figure 6. Learning is still completely characterized by the singular value decomposition, but now with a much stronger dependence of the strength of the singular values: The rate of acquisition of each dimension is now proportional to the square of the singular value as reflected in the exponential terms in the revised learning dynamics equation below:

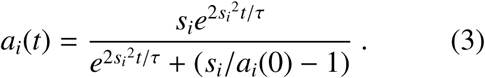

Thus, for example, a dimension 5 times stronger than another would be learned in 1/25 of the time required to learn the weaker dimension. This is reflected in the fact that the learning curves for the stronger dimensions are much steeper in the auto-associative case than in the case with one hot inputs, as can be seen by comparing to top and bottom panels of Figure 6.

In summary, while details of the dynamics of learning depend on details of the neural network architecture, training patterns, and the formulation of the learning task, the general characteristics of the time course of learning we observed with the analytically tractable deep linear network are conserved under a wide range of variations, supporting the view that the patterns of learning we have observed in our deep linear network are relevant to understanding learning in a wider range of cases. We therefore now return to the simple and analytically tractable deep linear case to explore its implications for learning new things in a network that has already learned the structure of a hierarchical data set.

### Learning Something New Building on Prior Experience

We now turn to applying our understanding of deep linear networks to the acquisition of a new item, once the content of a set of experiences has been learned. We focus on the case of the network with one hot inputs as shown in Figure 5, using the previously unused input unit shown in the figure for the new item. We have chosen an item that clearly belongs with other items already known within a specific category – in our case, the category of birds – but that, like the penguin in our earlier explorations, is not completely compatible with any specific already known bird. In particular, we consider an item we call the **sparrowhawk**, shown in Figure 7A. It has one of the features of the hawk (fierce) and one of the sparrow (small), and it has its own unique distinguishing feature, as all of the items in the environment do.

To begin our analysis, we apply singular value decomposition to the new complete data set, as shown in Figure 7B. Comparing this with Figure 4B, we see that the SVD of the new data set is similar to, but both alters and extends, the SVD of the original data set. In particular, the first and third dimensions of the original data set have been slightly altered, and a new dimension has been added (direct comparisons of differences between the SVD’s can be found in Figure 10, where they are linked to dynamics we will explore below). The first dimension now reflects the altered overall feature probabilities for the animals and the third dimension reflects adjustments to these probabilities needed to capture the average properties of the full set of birds and fish. Dimensions 1 and 3 are now a bit stronger than before, due to the added item participating in them, a fact most easily seen in Figure 10. The new dimension reflects how the individual bird representations must be adjusted to compensate for the adjustments to the other dimensions and to accommodate the sparrowhawk together with the existing birds accurately and without error. (By convention, the software that computes the SVD arranges the dimensions in descending order of strength, but we have rearranged them so that the sparrowhawk, and the new dimension associated with it, remain at the bottom in the diagram to make comparisons of other dimensions across the two data sets easier). The inset in Figure 7 shows the actual singular dimension strengths, indicating that the new dimension is somewhat stronger than the existing weak dimensions that separate the individual trees and the individual plants from each other. These dimensions are weak because these items differ only by a single item-specific differentiating feature.

It will be useful first to consider what happens if we now train a network we have previously trained on the eight original items, using focused training of the new item – that is, presenting the new item repeatedly without any interleaving with other items. As before, time in the simulation is represented in normalized time units corresponding to the number of epochs times the learning rate. It is crucial to understand that an epoch consists of a sweep through the training set now in use, which in the current case is just one item, the sparrowhawk.

The results of this simulation are illustrated in the top row of panels of Figure 8. In this and subsequent figures, we show separately the dynamics of the network error when tested on the new item (the sparrowhawk, leftmost panel), along with the dynamics of any error occurring on the other types of already known items (next three panels). In these panels, we have inverted the vertical axis so that progress in learning is reflected by an upward trajectory while interference results in a downward trajectory. The fifth panel shows the strengths of the dimensions of a singular value decomposition of the network’s output with all nine items in the full 9-item data set, measured with learning turned off so that we can see what the network knows without changing it, and the sixth panel shows the total summed error across all 9 items (without inverting the vertical axis, in accord with standard conventions in reporting this measure). The top left panel of the figure shows that the network reduces its squared error on the sparrowhawk to about 2.5 in a small fraction of a normalized time unit, corresponding to only a handful of presentations with a moderate learning rate. This outcome occurs because the sparrowhawk projects strongly onto the first and third existing dimensions of the data set, and much of the information about the sparrowhawk is captured quickly as the network exploits these projections. Concretely, this rapid improvement occurs because the output weights of the network already contain a dimension that captures what birds have in common with all animals and another that captures how the birds differ from the known fish, and the sparrowhawk projects strongly onto these dimensions, as shown in the SVD in Figure 7. Thus, the network quickly learns to map the sparrowhawk onto these two dimensions, capturing how it shares the average properties of the known birds, without any adjustment of the connection weights between the hidden and the output level – the existing weights from the hidden to the output units strongly propagate learning signals to the weights from the sparrowhawk input unit to the hidden units so that these input weights come to capture this average pattern very quickly, and these adjustments alone can reduce the sum squared error for the sparrowhawk from its original value of 9 (below the range of the four left panels in Figure 8) to 2.5. This point is demonstrated in Figure 9, where we have frozen the output weights (that is, prevented them from changing) for the first normalized time unit, so that all of the learning is occurring in the weights from the sparrowhawk input unit to the hidden units. The remaining discrepancies at the output level require adjustments to the hidden-to-output weights, to allow the network to reproduce the sparrowhawk’s output features exactly. This, however, produces considerable interference with the existing birds and, to a smaller extent, with the existing fish, since these items also project onto the same dimensions. Essentially, when tested with the existing birds or fish, the output will be distorted in the direction of the sparrowhawk. Critically, however, no interference occurs with the network’s knowledge of the four existing plants; performance on these items remains completely unaffected since the sparrowhawk is completely orthogonal to all of these items.

**Figure 8.**
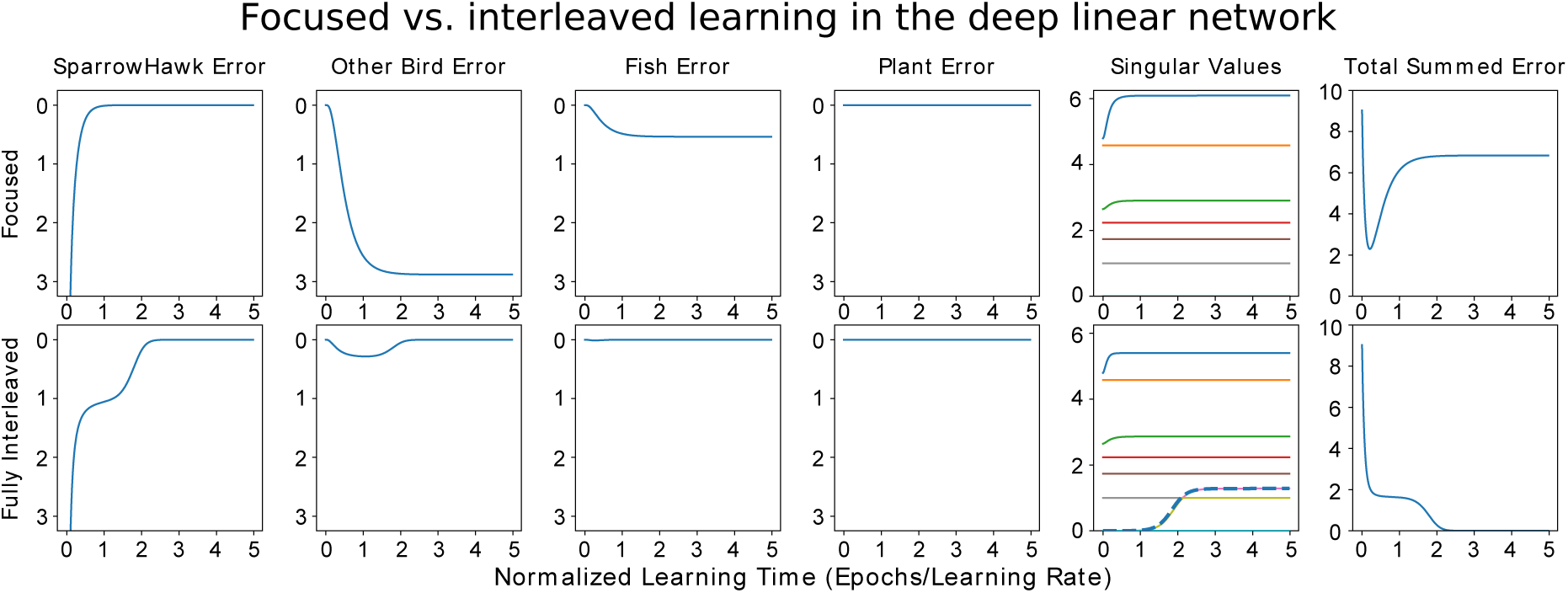
Focused vs. Interleaved Learning of a new item in the deep linear network with one-hot inputs, after prior learning of the eight-item data set. Top row: Focused learning condition: Only the new sparrowhawk item is presented in each epoch of training. The sparrowhawk is learned rapidly (left panel; note that the initial error on this item is 9.0, off of the scale shown; much of this error is eliminated within a small fraction of the first time unit), with severe interference with each of the known birds, milder interference with the known fish, and no interference with the known plants (next three panels). Each of the first four panels shows the average per item error for the set of items considered in each panel (1 sparrowhawk, 2 other birds, 2 fish, 4 plants). The fifth panel shows strengths of the singular dimensions in the network’s output as they change over the course of learning. Existing dimensions are adjusted, resulting in the observed interference. The sixth panel shows the total summed squared error over all nine items. Bottom row: Full interleaved learning: Here the sparrowhawk and all other items are presented once per training epoch. Learning of the sparrowhawk is retarded and there is only mild interference with the other birds. This is resolved when the network learns the new dimension that contrasts the sparrowhawk with the other birds. The emergence of this new dimension is shown in the fifth panel, where we see tat its strength begins to rise from 0 after about 1.5 time units. Note that the empirically measured learning curve for the new dimension (a solid line) coincides with and is largely hidden by the mathematically calculated learning curve for this dimension (solid line) when it is learned in a situation where all nine items are learned in a fully interleaved fashion without any prior learning.

**Figure 9.**
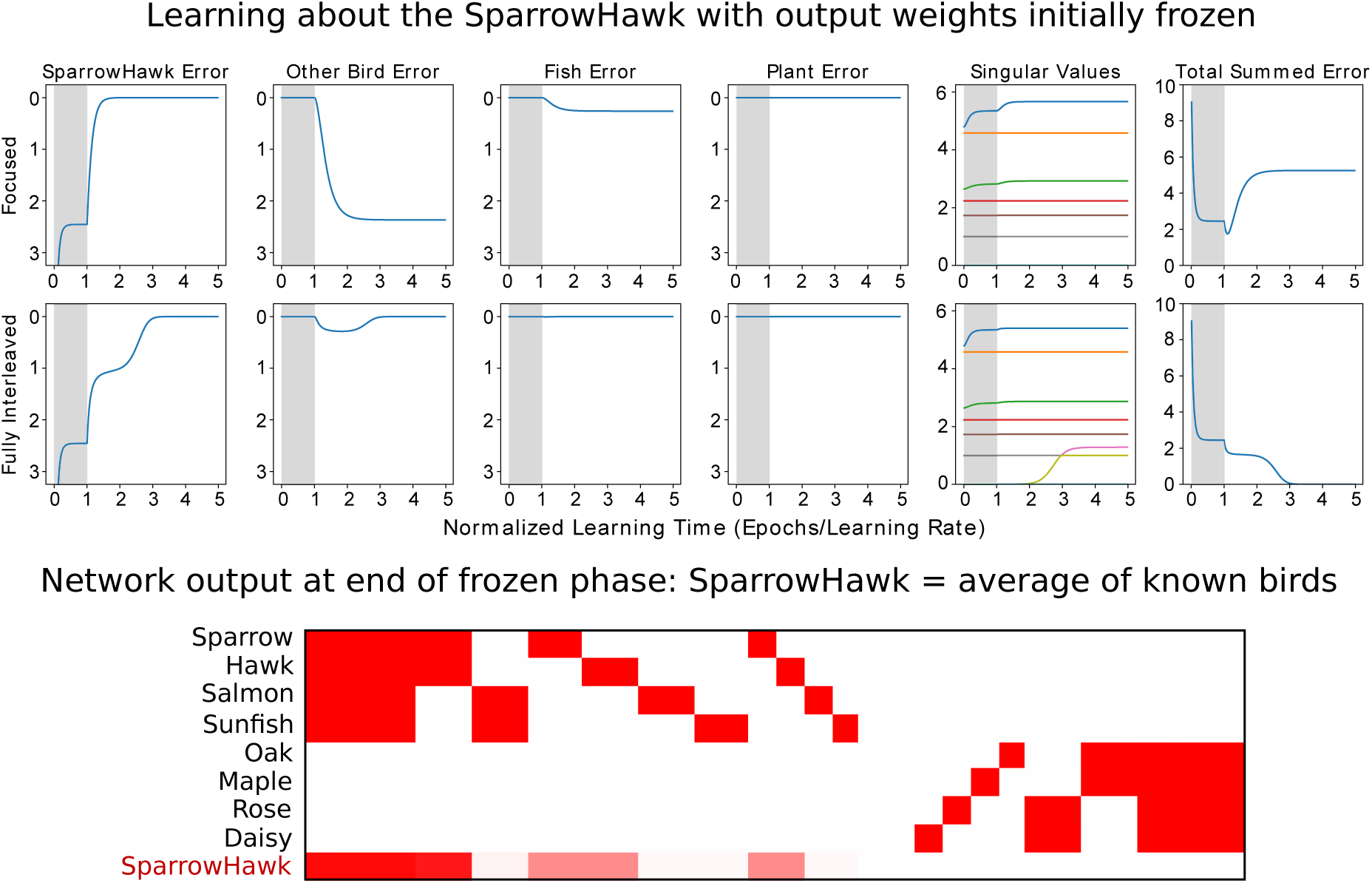
What the network can learn quickly about the sparrowhawk, based on the existing knowledge in its output weights. The top two rows of the figure show focused and interleaved learning results, as in Figure 8, with the change that for the first normalized time unit (shaded grey), the output weights of the network were frozen, so that all new learning occurred in the input-to-hidden weights. The bottom panel shows the network output for each input pattern at the end of the frozen phase. Because the sparrowhawk projects onto the known animal and bird-fish dimensions, it can quickly learn to map the new item onto these dimensions, even with the network’s output weights frozen, producing an output corresponding to the average of the known birds, as shown in the last row of the bottom panel. There is no interference with any of the existing items (first eight rows of bottom panel), since there is no overlap in the input weights among any of the items, and the output weights are prevented from changing. Once the output weights are unfrozen, learning proceeds largely as in the case where the weights were never frozen.

In summary, the network has learned about the new item by adjusting existing dimensions encoded in its connection weights. The essential problem with this is that, to fully accommodate the sparrowhawk together with the existing birds and fish, the network must learn the new dimension shown in Figure 7b. The network can learn the new dimension, accommodating all of the items perfectly, if it is trained using interleaved learning, as we now discuss.

Specifically, we now consider what happens if we continue learning from where we left off with the original eight items, but now with the sparrowhawk as an added, ninth item, so that one epoch now corresponds to the presentation of all nine items. The results of this simulation are shown in the bottom row of panels of Figure 8, and detailed visualizations of the changed and new dimensions are provided in Figure 10. For several of the dimensions, nothing has changed, and their initial values already correspond to the full values of *s*_*i*_. Therefore, no further adjustments occur to these dimensions. For dimensions 1 and 3, the projections are quite strong, and so the starting point is already quite far along the eventual learning curve. Learning thus proceeds immediately for these dimensions, as it did in the case of focused training with the sparrowhawk. This corresponds to the fact that aspects of the new item that are consistent with what is already known can be rapidly assimilated. As before, within a fraction of a normalized time unit, the aspects of the sparrowhawk that map onto dimensions 1 and 3 are learned. Over the rest of the first half of the first time unit, the network finds weights that represent a compromise of the first and third dimensions so that the other aspects of the sparrowhawk are partially accommodated (reducing the error on the sparrowhawk to about 1, as shown in the leftmost panel of the bottom row of Figure 8) at some small cost to the existing birds (as shown in the second panel of the bottom row). Only later, well into the second normalized time unit, does the network begin to learn the new dimension, so that it is fully assimilated partway through the third time unit.

**Figure 10.**
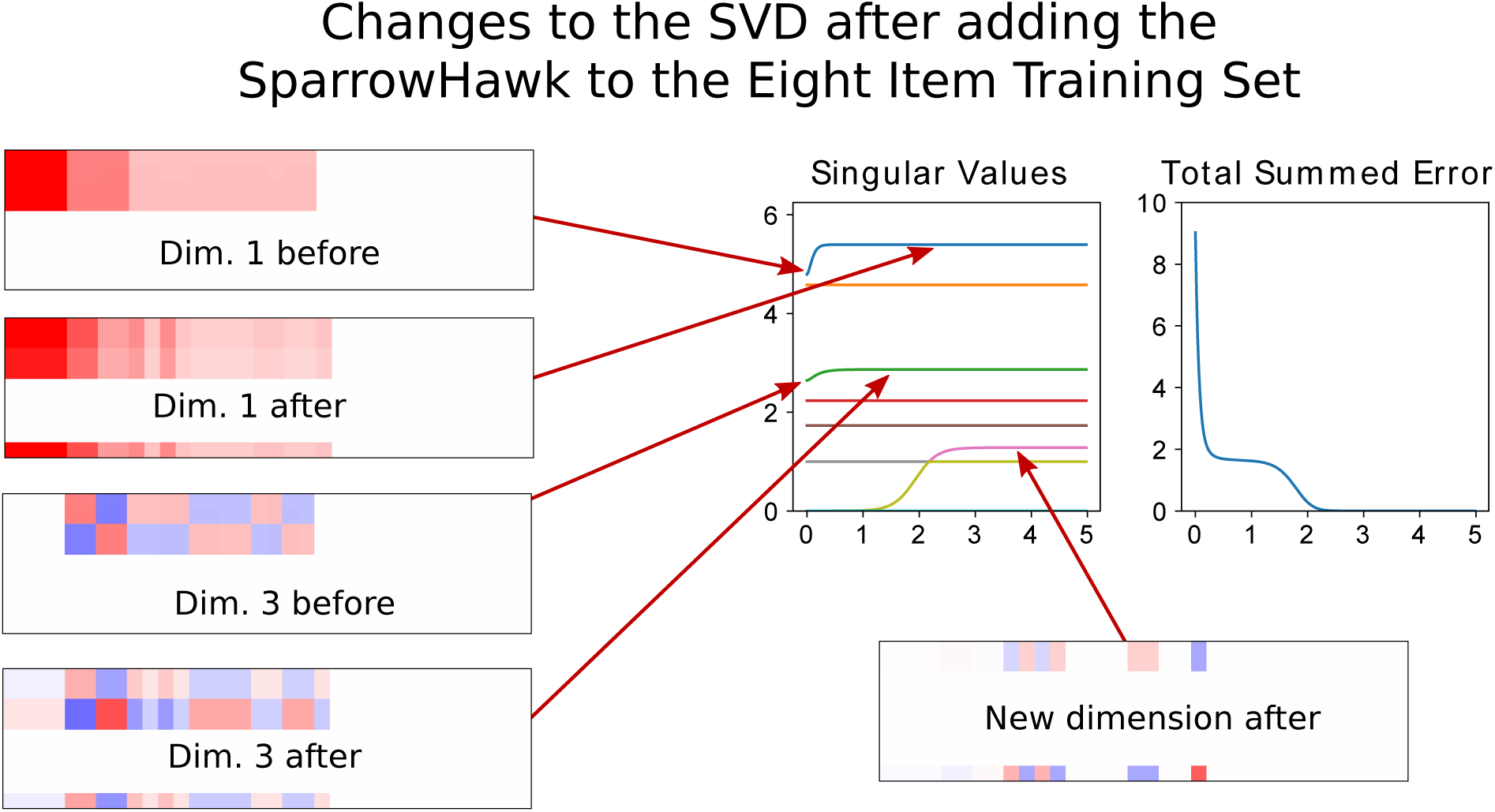
Detailed analysis of the changes in the SVD of the network’s output from the addition of the sparrowhawk to the eight-item training set, when trained with full interleaved learning. The first sion rapidly comes to reflect the new overall probabilities of animal features, and the third rapidly reflects the adjustments needed to capture the new overall probabilities of bird features. The new dimension that accommodates the sparrowhawk along with the already known birds is acquired after a delay. During this delay, the network’s output for the three birds reflects a compromise between their properties, and this constitutes interference with aspects of the sparrow and the hawk that contrast with aspects of the sparrowhawk.

Importantly, we find that the time it takes to learn the new dimension that fully allows all three birds to be effectively represented without compromise is just as long as the time it would have taken to learn this dimension had the network learned the whole data set all at once from scratch. This is documented in Figure 8, row 2, panel 5, which shows that the learning curve for this dimension as calculated using Equation 2 (shown as a dashed blue line) corresponds very closely to the observed learning time for this dimension (plotted as a solid line coinciding with and therefore hidden by the dashed line).

These observations have important implications for our general understanding of new learning of items in familiar domains. In one sense, the sparrowhawk is highly schema consistent, to use the terminology of Tse *et al.* That is, it shares all of the properties of animals in general and all of the shared properties of birds, and even has two properties that have already occurred in other birds. It is not, however, fully predictable from knowing that it is a bird. It has one unique property, and it combines variable properties found within birds differently than they have previously been combined. In general, this situation must apply to nearly every new thing we learn about. Something new will generally share some properties with previously known items, but some of its properties are likely to be unpredictable. Thus, in general, new items will have properties that are consistent with prior-knowledge and others that are not, and, at least in the present model, the aspects of a new item that can be assimilated easily are only those that are consistent with what is already known.

Before continuing, we consider both specific and general implications of the finding that the aspects of a new item that it shares with other items may be easier to learn than its more idiosyncratic aspects. Specifically, let us apply this observation to a consideration of the findings of Tse et al. [11, 12]. They found rapid neocortical consolidation of schema-consistent information in that their animals learned to map a new flavor onto a place in a familiar environment. Importantly, we note that the places animals had to associate with these new flavors were immediately adjacent to places already associated with previously learned flavors. For all practical purposes the animals could have mapped the new flavors exactly onto the familiar adjacent places since they would have effectively reached the new places by navigating to the already familiar ones. Thus, only the very easy part of neocortical consolidation may have been necessary in their study. Further research is needed to understand whether neocortical consolidation of a new location within an existing environment would occur equally rapidly; it could be that locations less similar with those already learned would require more gradual, interleaved learning.

More generally, the observation that some aspects of a new item can be integrated into a deep network quickly and without interference while others require extensive interleaving is worth exploring in a wider range of paradigms. We will consider empirical evidence related to this issue as well as possible future experimental tests in the general discussion.

### Learning Arbitrary Aspects of New Things Efficiently

We now turn our attention to a set of issues that go to the heart of questions about the role of experience replay in neocortical consolidation, starting from an issue that we raised in [7]. There we considered the fact that integration of new knowledge without interference into a neocortex-like network required interleaving with items previously learned, and we have just illustrated this point again in our new simulations with the sparrowhawk. We have refined the analysis to make clear that some aspects of information about a new item can be integrated quickly and without interference, but to fully integrate the new item may require extensive interleaved learning. A pressing problem then arises: Interleaved learning in all of our existing simulations has involved interleaving the new item with the full corpus of other items previously learned. This could be impractical at least for adult humans, since the totality of human knowledge would be very extensive indeed. This raises the question whether it is really necessary to be re-exposed to all of prior experience when learning something new, or whether, instead, more selective re-exposure to specific relevant experiences would be sufficient.

Our explorations with the sparrowhawk already hint at this possibility, since we have observed that the interference that occurs with focused learning of the sparrowhawk is greatest for the existing birds, only moderate for the existing fish, and absent for the known trees and flowers, as shown in the top row of panels of Figure 8. We therefore considered whether focusing interleaving on the items that are similar to the new to-be-learned item could allow integration of knowledge of the sparrowhawk without requiring full interleaving with all previously learned items.

Accordingly, we conducted additional simulations. In one of these, we employed *similarity weighted interleaved learning* (SWIL). Here, each training epoch involved 1 presentation of the sparrowhawk and the previously known birds, .2 presentations of the previously known fish, and no presentations of any of the trees or flowers, leading to a total presentation rate of 3.4 items per epoch.^5^ In the other *uniform interleaving* condition, the new sparrowhawk item was also presented once per epoch, but now the other 8 items were each presented at a uniform rate of .3 presentations per epoch, resulting in the same total presentation rate of 3.4 items per epoch. The results, shown in Figure 11, indicate that the similarity weighted regime results in virtually identical results on a per epoch basis compared to full interleaving, but each epoch now includes less than 40% as many pattern presentations. Thus, we can achieve the same results we obtained with full interleaving, using 2.5 times fewer presentations. In contrast, the uniform interleaving condition is far less efficient, slowing down the acquisition of the new dimension required for full integration of the sparrowhawk with existing knowledge. The amount of slowdown corresponds nearly exactly to the extent of the reduction in exposure to the relevant patterns (the three birds). This correspondence is indicated by the coincidence of the observed learning time for the new dimension, indicated by the solid curve rising from 0 starting part-way through the third normalized time interval in the singular value plot for the Control condition, and the learning time that would be expected due simply to the reduced exposure to the three birds, indicated by the dashed curve. The dashed curve was obtained by scaling the theoretical learning curve for the new dimension already shown in Figure 8 by the ratio of the average bird presentation rate (1.0) in the full and similarity weighted conditions to the bird presentation rate in the control condition ((1 + 2*.3)/3, or .5333). Essentially, then, the control condition just slows learning down proportionally to the amount of exposure to the patterns contributing to the new, to be learned dimension, whereas the SWIL condition results in full integration without unnecessary exposure to completely unrelated aspects of existing knowledge.

**Figure 11.**
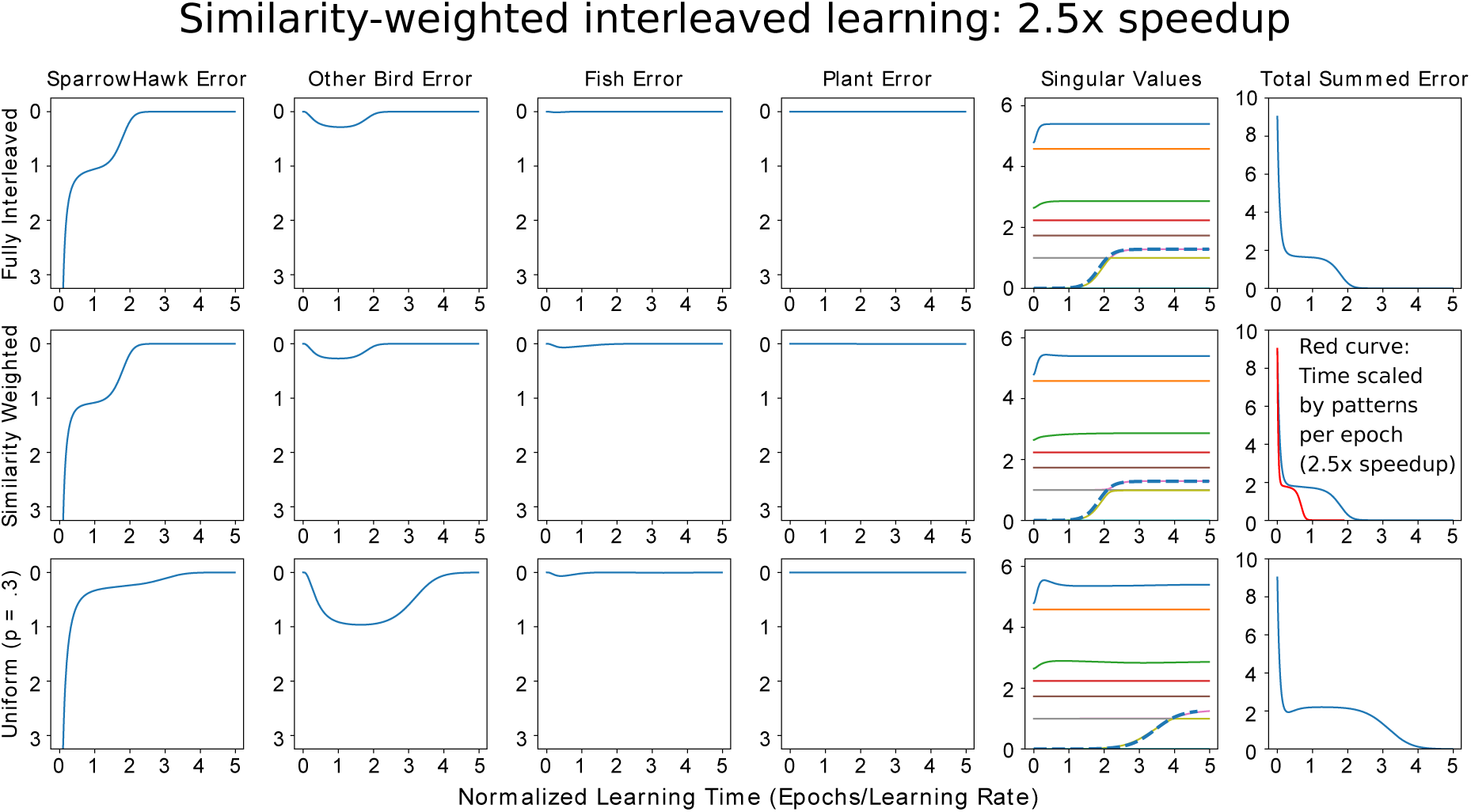
Similarity weighted interleaved learning (middle) compared with full interleaving (top, reprinted from Fig 8 for comparison) and a uniformly weighted control condition in which each known item is presented .3 times per epoch, equating the number of patterns per epoch with the similarity-weighted condition. Average number of pattern presentations per epoch is 9 for full interleaving, and 3.4 for the other two cases, or 38% of the pattern presentations with full interleaving. Dashed lines show expected learning time for the dimension that differentiates the sparrowhawk from the other birds when all items are learned in a fully interleaved fashion without any prior learning (top two rows), or when scaled for the average number of presentations of relevant items (i.e. the birds) in the control condition (bottom row). In the rightmost panel, the value of the squared error measure is plotted as a function of normalized learning time per epoch in blue. The curve in red in the middle row scales the blue curve by the number of patterns per epoch in the similarity-weighted condition to show the 2.5 x reduction in total pattern presentations that resulting from similarity weighted interleaved learning.

### New Learning in the Deep Linear Auto-Associator

As we have previously discussed, our analysis of the dynamics of learning in deep linear networks also extends to the deep linear auto-associator, a network architecture that is interesting for reasons considered earlier. We already saw how gradual learning within a deep linear auto-associator results in the gradual differentiation of representations at different hierarchical levels. In Supplementary Materials, Section II, we consider the process of acquiring a new distinct representation corresponding to the sparrowhawk in a deep linear auto-associative network. These simulations demonstrate corresponding patterns of learning and interference over the same range of regimes considered in the pattern associator case. The main difference is that, after learning the eight patterns in the base training set, the network already knows how to capture most of the content of the sparrowhawk pattern. Presenting the sparrowhawk pattern on the input to the auto-associator after it has been trained on the original eight item training set results in the output pattern corresponding to the average of the existing birds (the same pattern that was produced in the pattern associator after learning about the sparrowhawk with frozen output weights, as shown in the last row at the bottom of Figure 9), giving rise to a sum squared error of 2.5 between the correct sparrowhawk output and the average bird pattern. This happens because the input features of the sparrowhawk project fully onto dimensions 1 and 3 of the knowledge already in the network, and these dimensions together capture the average properties of the existing birds. As before, to integrate the sparrowhawk into the network’s weights, a new dimension must be learned. Focused training with the sparrowhawk alone produces interference with the existing birds, while either full or similarity weighted interleaving, as in the pattern associator case, results in the gradual acquisition of the new dimension that accommodates the sparrowhawk along with the other previously-known birds.

### Pre-Training Effects in Deep Neural Networks

The observations we have made in the above sections allow us to relate our analysis to some issues that have been raised in previous research on the role of auto-associative pre-training in deep neural networks. Such pre-training is often called unsupervised pre-training, because only the items, without labels, are used during pre-training. Prior research (e.g., [29]) has demonstrated that such pre-training of a deep non-linear neural network speeds later learning of an input-output mapping, and the theory of deep linear networks has been applied to this case [30]. In terms of our example, consider what would happen after training our linear auto-associator with our eight item data set if we then added a new output layer containing a one hot classification unit for each of the eight items. If we now train this network to map from the input to these one-hot output units, the input-to-hidden weights would already capture all of the relevant dimensions of the input, and so it would only be necessary to adjust the output weights to learn to classify each item correctly. This point was demonstrated in simulations with the MNIST classification data set in [30], which also showed that the properties of deep linear networks that we have reviewed here are largely conserved regardless of the number of layers in the neural network. Extrapolating from the findings reported in [30], we expect that learning to classify the items in our simpler plant-and-animal data set would occur very quickly and would not require any change to the input weights *as long as the pre-training spans all of the relevant dimensions*. Note that pre-training on the original eight items would result in a representation that does not span the new dimension required to learn to accommodate the sparrowhawk. More gradual, interleaved learning and adjustment of the input-to-hidden weights would be required to learn the added dimension needed to correctly identify this item without interference or confusion with the other birds, just as it has been in the simulations we have reported above.

We hope that this brief section helps to suggest the relevance of the theory of deep linear networks to a wider range of deep neural network applications for both machine learning and computational neuroscience. The theory does not cover all aspects of the phenomena observed in networks with non-linearities, a point we return to under *Future Directions* below.

## General Discussion

### Summary and Implications for Complementary Learning Systems Theory

In the preceding sections of this article, we have observed several aspects of learning in simulated deep neural networks, and we have offered a formal theory of learning in deep linear networks that captures and extends many observations made in earlier publications. We view these networks as capturing aspects of cognitive and conceptual development explored in more detail elsewhere [20], and as providing a framework that has helped inspire our initial investigation of complementary learning systems [7]. Throughout this work, we have used deep networks to model the structure-sensitive learning process that appears to take place within the neocortex, helping us to understand what the complementary, hippocampus-dependent learning system must do to complement this structure sensitive learning system.

A central outcome of our earlier work was the observation that deep neural networks learn to capture the structure in a domain of experiences in a gradual, progressive, and stage-like fashion. While they could learn new things quickly by making large connection weight changes, such large changes would lead to interference with existing knowledge. The hippocampus-dependent system complements this system by supporting rapid learning of new knowledge without interference. This new knowledge can then be gradually integrated into neocortical structures through interleaved learning with ongoing exposure to items that had previously been learned.

Challenged by the important findings of Tse *et al.* [11, 12], and with the benefit of further simulations, an important qualification of these points was introduced in [18]. There it was noted that new knowledge highly consistent with what is already known can be integrated rapidly into neocortex-like deep neural networks, providing a start toward a computational understanding of rapid neocortical consolidation of schema consistent knowledge, as demonstrated by Tse *et al.*. The work presented here has further extended this work, leading to several additional key observations.

Perhaps the most important new observation is that different aspects of experiences might not always be equally easy or hard to learn, either by biological learners or deep neural networks. The sparrowhawk example we have considered here exemplifies properties of many of the new things that we learn about. New things often share properties with things we already know, but like the sparrowhawk, they are not fully predictable from those prior experiences in two ways. New items can have their own idiosyncratic content, represented by unique features not shared with other items, and they may recombine aspects of known items in novel ways. New types of things (like a new species of bird) or new individual persons, places, or things, generally have many properties that are typical of their super-ordinate category, while also having unique properties or combinations of properties. Though our observations with the sparrowhawk were implicit in earlier work, the present work has brought out more clearly that it may be possible to learn some aspects of new things very quickly and without interference. It isn’t *everything* about something new that is hard to learn in a deep neural network, it is *the aspects of it that di*ff*er from other things* that can be hard to learn.

Another important observation is that the interference with existing knowledge that arises from learning something new is also not completely general. As we have seen, new learning about the sparrowhawk interferes only with knowledge of similar things, and even then, the interference is restricted to those aspects of the similar items that are in conflict. There is no interference with knowledge of completely unrelated things at all, and no interference with knowledge of aspects of similar items that the new item shares with others it is similar to.

These observations are captured in our simulations with a simple hierarchically structured dataset learned by a simple version of a deep neural network. Our analysis, building on [26], reveals that learning in these networks must be understood in terms of learning the *structure* in the ensemble of items from which the network learns, rather than simply in terms of such matters as the frequency of exposure to the individual items themselves. The dimensional structure of the ensemble of items – as captured by the singular value decomposition – dictates the learning time needed to acquire knowledge, both when that information is being learned from scratch, and when it is being added to a body of knowledge already known. We can rapidly integrate the projection of the sparrowhawk onto existing knowledge structures. However, the novel features and novel combinations of features require a new dimension to be integrated into the structured representation in the weights of the network and thus require interleaved learning if they are to be learned without interference with other similar items.

It is important to acknowledge that the networks we have focused on here are far less complex than the real neural networks in the brain. We have analyzed networks that are completely linear and contain only two layers of modifiable connection weights, and the neural networks in the brain surely contain more layers and exploit non-linear computations. While our simple networks capture many properties of learning in the deeper, nonlinear network we used in previous work, greater depth and non-linearities add computational capabilities that our networks do not have. Thus it is important to be cautious in extrapolating what we have learned to deeper and more non-linear networks or real biological systems, and further work is needed to explore the limits of any such extrapolation. Nevertheless, it seems worthwhile to consider the implications of our findings for understanding the roles of replay and interleaved learning in our understanding of the neural basis of learning and memory.

### Implications for Replay and Interleaved Learning

To our knowledge, the idea of memory replay originated in the theoretical speculations of Marr [8], who proposed that the hippocampus stores on the order of 10,000 experiences every day, and replays them overnight to allow the cortex to sort them into categories and to adjust these category representations. Inspired by this, Wilson and McNaughton [25], building on Pavlides and Winson [31], were able to demonstrate that correlated patterns of neural activity occurring during waking behavior were subsequently recapitulated during subsequent sleep episodes. Our extension of Marr’s ideas, as elaborated first in [7], offers a different take on the role of replay, however. In our theory, it is not just new information that needs to be re-experienced for integration into neocortical structures. Instead, the integration of newly encountered items or experiences with unique properties or novel combinations of properties requires interleaving with existing knowledge if this integration is to occur without interference.

One view worth keeping under consideration is that much of the experience that is required for integration of new knowledge may come from ongoing experience. For example, if a new pet enters our life, we would experience it every day, and as we continue to go about our daily routine, we would continue to be exposed to many other things. The hippocampus could play an important role in initial formation of a memory for the new pet. Some aspects consistent with existing knowledge would be easy for the cortex to learn, while others, such as remembering the name of a new pet or its particular proclivities would remain hippocampus dependent for a longer period of time. Gradually the cortex would learn through repeated exposure, so that the hippocampus would no longer be necessary. Meanwhile we would have ongoing exposure to other things including other people’s pets and other animals, avoiding the problem of catastrophic interference. A question arises here: What, on this view, would be the role of replay of new information during off-line periods shortly after initial exposure to new information? While this replay is often thought to begin the process of integration into the neocortex, another important possible role would be to help stabilize the plastic changes within the fast-learning hippocampal system, ensuring that the new learning remains available for use and replay over an extended time period. If such replay events were selective for repeated and/or highly salient aspects of our recent experience, they would help conserve limited synaptic resources in both hippocampal and neocortical circuits, reserving them for material likely to be of use in the future.

We have also explored a factor that could potentially speed the integration of arbitrary new information into neocortex, namely hippocampus-dependent replay of recent experience interleaved with selective replay of similar already known information. We observed that learning arbitrary new aspects of things without interference with existing knowledge does not require ongoing exposure to all existing knowledge; instead it can be enough to focus only on related items of information, relying on an experience protocol we have called similarity-weighted interleaved learning. Similarity-based reactivation of pre-existing knowledge could occur, in part, during direct experience with new things. Returning to the example of a new pet, when we have an experience with it, that experience may trigger memories of other pets – our own or those of our friends and acquaintances. Thus, it is possible that similarity weighted exposure to related things we already know about may be a natural concomitant of experience with new things.

While ongoing experience may play a role in promoting interleaved learning, some studies support the idea that replay without ongoing exposure can also lead to neocortical consolidation. In one study showing this pattern, Kim and Fanselow [32] exposed rats to highly aversive tone-shock pairings in a single session in a novel environment. They found that removal of the hippocampus one day after the experience resulted in little or no expression of fear when the animals were returned to the environment, but leaving the hippocampus intact for a period of one to four weeks after this experience resulted in gradually increasing expression of fear after a subsequent hippocampal lesion. The increase in fear memory occurred while the animals were retained in their home cages, with no re-exposure to the fear-inducing environment. While there are many studies that have failed to show increasing retention after a longer period prior to a hippocampal lesion [33], there are other sources of evidence of gradual integration into the neocortex. For example, Takehara-Nishiuchi and McNaughton [34] demonstrated the emergence of neural activity in deep layers of the medial prefrontal cortex over a a period of several weeks without ongoing task exposure, also supporting the idea of gradual integration of initially hippocampus-dependent learning into non-HC-dependent structures through off-line replay.

In the context of the evidence for gradual integration of new knowledge into the cortex without ongoing exposure from the environment, it is intriguing to consider the possibility that similarity weighted interleaved learning might occur during off-line periods based on information already stored in the brain. In ongoing work, we are exploring the possibility that a hippocampusdependent replay of a recent new memory would activate the corresponding representation in the neocortex, providing a learning trial for the cortex based on the new memory. The initial cortical activation during the novel experience might also leave a residual trace within the cortex that may bias the cortex, so that spontaneous neural activity would tend to reactivate memories stored in the neocortical connection weights with neural activity patterns that overlap with the one that represents the new experience. In future work, we hope to explore this possibility through computer simulations relying on more biologically grounded simulated neural networks than the ones we have employed in the work presented here.

### Open Questions for Future Research

While the work reported here demonstrates that similarity weighted interleaving can allow new information to be fully integrated with less ongoing exposure to existing knowledge than might otherwise have been thought, we have not found a similar advantage for similarity weighting with some other data sets and neural network architectures. In ongoing work we are exploring this issue. It should be clear from our simulations that degree of overlap with other patterns is a relevant factor that influences the efficacy of similarity weighting, and future work should explore whether sparse coding schemes that reduce overall pattern similarity might enhance the advantage of similarity weighting. Perhaps, if each pattern to be learned only overlaps with a small fraction of the other patterns, then interleaving might only be required for the items in this small set. On the other hand, excessive sparsity can reduce a network’s ability to capture relevant similarity structure and thereby to generalize in useful ways, and cortical networks may therefore exploit an intermediate level of sparsity that balances the need to capture generalizations while minimizing interference [35]. In summary, the extent of the advantage of similarity weighting is likely to depend on the details of the similarity relationships among the patterns to be learned, and future work is needed to explore more fully the advantages that similarity weighting might provide in a wider range of situations and to explore how the representations of items to be learned might be tailored to promote the effectiveness of similarity weighting.

More broadly, it should be noted that our work does not fully address the broader literature on learning structured bodies of knowledge and the roles of schema consistency and inconsistency in new learning. In particular, much of the work in humans highlights an important role for novelty as well as schema consistency in driving memory replay, and a role for the medial pre-frontal cortex in this process [14], and other evidence suggests that less consistent information may be prioritized for replay [36, 17]. A complete theory of new learning will require a fuller understanding of these important aspects of the memory formation and replay process [37].

### The Role of Models in Cognitive and Systems Neuroscience

Before closing, a comment may be in order about the relationship between computer simulations and experimental investigations of real neural systems. The role of models is not to capture reality in all of its complexity, but to simplify it so that some of its key properties may be understood [38, 39]. The simple simulations we present here do not fully capture what really happens in the brain during learning. We strongly believe that further progress will still require simplification, but that any consideration of what is observed in the behavior of simulation models must also be viewed as providing a limited perspective that needs to be complemented by consideration of the properties of the real neural system. It is only though such interplay that real progress will be made.

## Conclusion

In spite of many years of progress in the psychology, neurobiology, and computational analysis of learning and memory, there remains a great deal we still have to learn. Real brains are far more complex than the simple networks we have considered here, and so is real experience. The discovery of the medial temporal lobe amnesic syndrome, the invention of methods that allow us to study neural activity during experience and during later offline periods, and the exploration of artificial neural networks that allow aspects of learning and memory to be captured in simulations have provided a starting place, and the findings from the research that has ensued in the wake of these developments constitutes real progress. Yet, they still leave us with only a partial understanding of how experiences and their replay contribute to learning and memory. We hope that the ideas and findings we have presented here will contribute to the ongoing exploration of these issues, and we expect that a full understanding will require ongoing research of all of the types we have mentioned for many years to come.

## Acknowledgments

This work was supported in part the Defense Advanced Research Projects Agency (DARPA) through grant HR0011-18-2-0021 (BLM) and contract FA8750-18-C-0103 (JLM). AKL was supported by an NSF Graduate Research Fellowship. A Hilgard Faculty Scholar Award from the Department of Psychology at Stanford University to BLM provided partial support for an extended visit during Autumn, 2017, facilitating conversations that gave rise to this research.

## Footnotes

1 The data set files and iPython notebooks that implement the singular value decomposition and deep linear neural network simulations described below and in the supplement can be found at https://github.com/lampinen/integration_CLS.

2 The quantities computed by PCA can be derived from the SVD and *vice versa*, if the column means are first subtracted from the item property matrix before calculating the SVD. In that case, the singular values in the SVD are proportional to the square roots of the eigenvalues in PCA. See [27] for more details. The quantities captured in the SVD without subtracting the column means are directly related to the dynamics of learning in our model, and so are more useful for present purposes

3 We follow the mathematical convention in which a bold lowercase letter such as **x** corresponds to a column vector and a bold uppercase letter such as **X** corresponds to a matrix. The superscript *T* stands for the transpose operation, which turns columns into rows and *vice-versa*. To better support intuition, we transpose the data matrix employed in [26]; this swaps the roles of the **u**^*i*^ and **v**^*iT*^.

4 The adequacy of the approximation depends on having enough hidden units to make it likely that there is a dimension embedded in the initial weights such that each to-be-learned dimension projects reasonably strongly onto it. A perfect match to the equation can be obtained by initializing the network with weights that exactly capture the to-be-learned dimensions scaled down to align with the chosen value of *a*_*i*_(0).

5 The fractional presentations of items are obtained by scaling down the learning rate. We also ran simulations in which fractional presentation rates are implemented by presenting items probabilitically. Those simulations produced results (not shown) that jittered around the smooth curves obtained by scaling down the learning rate.

