## Supplementary Materials for "Integration of New Information in Memory: New Insights from a Complementary Learning Systems Perspective"

This Supplement contains two sections:

I. Some theoretical and empirical investigations of mode adjustment in deep linear networks.

II. The dynamics of learning a new item in a pretrained autoassociative network.

### I. Some Theoretical and Empirical Investigations of Mode Adjustment in Deep Linear Networks[1]

#### Mode Learning Dynamics

Saxe et al. [1] provided (in part A of their supplementary material) solutions to learning dynamics in linear networks from arbitrary initial mode strengths, with some assumptions about the initialization and the structure of the data. We note a minor correction to these solutions (see below) and build off them here.

Consider the input-output correlation matrix $\Sigma_{31} = \sum_{i=1}^{P} y_i x_i^t$ where $\{(x_1, y_)), ..., (x_P, y_P)\}$ are the (input, target) pairs the network is trained on. Saxe and colleagues considered its singular value decomposition:

$$\Sigma_{31} = \sum_{\alpha=1}^{k} u_\alpha s_\alpha v_\alpha^T$$

Saxe and colleagues assumed that the input-input correlation matrix is white ($\Sigma_1 1 = \sum_{i=1}^{P} x_i x_i^t = I$), and under the assumption that the network is initialized so that the singular value modes are decoupled, they showed that the modes then remained decoupled and gave exact solutions for the learning of these modes from small initial weights. In the supplementary material, they also expanded this to arbitrary initial weight size (but still assuming decoupled initialization).

Specifically, consider singular mode $i$. For ease of explanation, we change the basis of the representational layer of the network so each mode is represented by a single hidden unit – this is permissable because we assumed the modes were decoupled. We call this the SVD basis. (This is equivalent to the change of variables denoted by bars by Saxe and colleagues.) Let the initial projection of this unit's input weights onto the input mode $v_i$ be $a(0)$, and the initial projection of its output weights onto the output mode $u_i$ be $b(0)$. Saxe and colleagues showed that $(a(t), b(t))$ evolve over time along hyperbolas of constant $a^2 - b^2$ until they approach $ab = s_i$, i.e. the true strength of that mode in the data.

[1]This section was developed and written by AKL.

We assume $a(0) \neq b(0)$ (otherwise a hyperbolic parameterization does not work). Without loss of generality we assume $a(0) + b(0) > 0$ (the other half-space requires a trivial reparameterization). We can then parameterize this hyperbola by the angle $\theta$ and make the change of variables

$$a = \sqrt{2c_0} \cosh \frac{\theta}{2}, \qquad b = \sqrt{2c_0} \sinh \frac{\theta}{2}$$

Where $c_0 = \frac{1}{2}(a(0)^2 - b(0)^2)$ so that

$$ab = c_0 \sinh \theta$$

Following the derivation of Saxe et al. [1] with this change of variables, and adding a factor of 2 that was omitted in their original derivation, we arrive at:

$$\frac{\tau}{2} \frac{d\theta}{dt} = s_i - c_0 \sinh \theta$$

(the factor of two can also be absorbed into the time constant $\tau$, we leave it separate here to avoid changing the definition of $\tau = 1/\lambda$ from the original paper).

This differential equation is separable, and so we can solve for the time needed to traverse along the hyperbola from an initial point $\theta_0$ to a final point $\theta_f$:

$$t = \frac{\tau}{\sqrt{c_0^2 + s_i^2}} \left[ \tanh^{-1} \left( \frac{c_0 + s_i \tanh\left(\frac{\theta}{2}\right)}{\sqrt{c_0^2 + s_i^2}} \right) \right]_{\theta_0}^{\theta_f} \tag{1}$$

This provides an exact analytic solution for the time a given degree of learning from a given starting point requires. Although this equation cannot be analytically inverted to find $\theta(t)$ (and thereby $a(t)$ and $b(t)$), we can parametrically sweep through the interval $(\theta_0, \theta_f)$ to plot the theoretical learning curve. In Fig. 1 we demonstrate the match between this theoretical learning curve and the empirical results for a linear two layer network learning a single mode, starting from a random initialization, which provides the initial $\theta_0$.

We note that in the special case that $s = 0$, for instance when a mode is removed from the data and is being unlearned, the solution is:

$$t = -\frac{\tau}{2c_0} \left[ \ln \tanh \left( \frac{\theta}{2} \right) \right]_{\theta_0}^{\theta_f} \tag{2}$$

We also note that

$$\frac{d(ab)}{dt} = a\frac{db}{dt} + b\frac{da}{dt} = 2c_0(s_i - c_0 \sinh\theta)\left(\cosh^2\frac{\theta}{2} + \sinh^2\frac{\theta}{2}\right)$$

Which corresponds to

$$\frac{d(ab)}{dt} = (s_i - ab)\left(a^2 + b^2\right) \tag{3}$$

That is, the change in the network's representation of a mode is proportional to the product of two factors: how far the projections are from the correct value (i.e. the error) and to how (absolutely) large the summed, squared alignments to the true input and output modes are. The sigmoidal trajectory of mode learning noted by Saxe and colleagues can easily be interpreted from this equation – with a small weight initialization, the second factor is small, and so the change in the strength of the mode is initially small. As $a$ and $b$ increase, the rate of change increases, until $ab$ approaches its asymptotic value of $s_i$, at which point learning slows down as the error shrinks.

Note that the theoretical results assume that the network's input and output modes are perfectly aligned with the data modes, and all that must be learned is the correct singular value. This assumption will not generally hold when new data are introduced after prior learning, because some modes will be adjusted. Nevertheless we find empirically that the theory provides decently accurate approximations even in this more general setting, as we show below.

**Losses.** The loss of the network at a given point in learning is given by a relatively simple formula. For singular dimension $i$ in the data let $\mathbf{v}_i^T$ be the input mode, $s_i$ the singular value, and $\mathbf{u}_i$ the output mode. Similarly, for each mode $j$ in the SVD of the outputs produced by the network, let let $\hat{\mathbf{v}}_j^T$ be the input mode, $\hat{s}_j$ the singular value, and $\hat{\mathbf{u}}_j$ the output mode

$$\text{Loss} = \sum_i s_i^2 + \sum_j \hat{s}_j^2 - 2\sum_i\sum_j s_i\hat{s}_j\left(\mathbf{u}^i \cdot \hat{\mathbf{u}}^{\mathbf{j}}\right)\left(\mathbf{v}^i \cdot \hat{\mathbf{v}}^{\mathbf{j}}\right)$$

For a derivation of this formula see Lampinen and Ganguli [2].

Although the formula above is more general, it is useful to consider the special case where each of the network's non-trivial modes has a non-zero projection onto only one of the data modes. This does not require that the network modes be perfectly aligned with the data modes, merely that the network modes be effectively "paired up" with the data modes so each only projects onto a single one. In this case, the loss per component can be calculated independently for each mode $i$:

$$\text{Loss}_i = s_i^2 + \hat{s}_i^2 - 2 s_i \hat{s}_i \left(\mathbf{u}^i \cdot \hat{\mathbf{u}}^{\mathbf{i}}\right)\left(\mathbf{v}^i \cdot \hat{\mathbf{v}}^{\mathbf{i}}\right) = s_i^2 + \hat{s}_i^2 - 2 s_i a_i b_i$$

where $a_i$ and $b_i$ are, respectively, the input and output mode projections as in the previous section.

#### Learning from Different Starting Points

We are now in a position to examine the question of how new knowledge gets integrated into linear networks that have already learned something. Given the above, this reduces to the question of how this new knowledge projects onto the knowledge that is already stored in the network. Qualitatively, new knowledge which provides a minor adjustment of existing knowledge will be rapidly integrated, since the projections of the new singular dimensions onto the old singular dimensions will be strong (i.e. the second factor of $d(ab)/dt$ will be large), and the first factor will be proportional to the amount of adjustment needed. Thus **adjustments to old modes will be rapidly integrated**, so long as they are not large enough to make the mode nearly orthogonal to the pre-adjustment mode. By contrast, entirely new knowledge (i.e. modes that are orthogonal to all previously learned modes) will be integrated quite slowly. In fact, it will be learned **over the same period of time as it would have taken to learn this mode in a randomly initialized network**, assuming the modes are decoupled.

In Fig. 2 we demonstrate a fairly close match between theoretical and empirical learning for these cases. In particular, the theoretical and empirical loss are very closely matched. However, the alignment of the mode being adjusted is slightly slower than the theory predicts. In fact, this is because there is a transient decrease in the singular value of this dimension while the network adjusts it, which is not predicted by the theory (since the theory assumed that the modes would be aligned, and only the singular values would differ). However, if we scale the alignment by the ratio $s/\hat{s}$, i.e. the ratio of the singular dimension strength in the data to the current network singular dimension strength, the empirical curve matches very closely here as well, see Fig. 2(b-c). Note that this is not a theoretically-derived adjustment, but rather an empirical observation that might lay the groundwork for future work towards a deeper theoretical understanding of the adjustment process.

Note that it suffices for only the input weights or the output weights but not both to have a strong projection onto the new mode for the adjustment to be rapid – slow initial learning only occurs when both projections are small, as in the case of small random initializations. This is clear from the expression for $d(ab)/dt$ given above.

#### Learning Multiple Non-Orthogonal New Modes

This theory assumes each mode is being adjusted (or learned from scratch) in a way that is orthogonal to all prior modes. However, often a new mode has projections onto an old mode, as in the case of the sparrowhawk. In fact, the adjustment made to the old mode is often precisely to remove its initial alignment to the new mode.

Fortunately, we have some understanding of what happens in these situations. When an adjusted mode has some projection onto two previous modes, the corresponding representational modes will *compete* over the adjusted mode. Similarly, when the adjusted mode and the new mode both share some projection onto the old mode, they will *compete* for its representational mode. The one with the strongest singular value and strongest projection onto the representational mode will win out and be learned first, all else being equal. However, this competition changes the representational modes and delays the incorporation of the new information. It is difficult to obtain exact analyses of learning in this situation, as the modes are no longer decoupled and their evolution can be quite complex. The overall pattern, however, is that **competition with a partially-aligned new mode will delay the adjustment of the old mode**, and this delay will be worse the more similar the strengths of the projections of the old modes onto the new mode are. This can be seen in Fig. 3 by the way the empirical learning curves lag behind the theoretical curves initially.

However, in this case the new mode will benefit slightly from the initial strong projection. Even though the mode being adjusted will win most of the representation and prevents the new mode from using these strong weights from before, the competition will actually end in a slight compromise, wherein the new mode will steal a little bit of this original representational mode away from the adjusted mode. This will result in a stronger earlier projection for the new mode. Because of the competition from the stronger mode being adjusted, the amount of this projection will be quite small in absolute terms. However, since the delay in initial learning of a mode is due to the lack of this early projection, the main effect of some projection of a new mode onto an old mode is a **slight acceleration in the learning of the new mode**. This can be seen in Fig. 3 by the way the empirical learning curves lead the theoretical curves late in learning.

#### II. New learning in the Deep Linear Auto-Associator

Here we describe details of the process of acquiring a new distinct representation corresponding to the sparrowhawk in a deep linear auto-associative network, demonstrating corresponding patterns of learning and interference over the same range of regimes considered in the pattern associator case. The results of these simulations are shown in Figures 4 and 5.

As noted in the main text, after learning the eight patterns in the base training set, the network already knows how to capture most of the content of the sparrowhawk pattern. Presenting the sparrowhawk pattern on the input to the auto-associator after is has been trained on the original eight item training set results in the output pattern corresponding to the average of the existing birds (the same pattern that was produced in the pattern associator after learning about the sparrowhawk with frozen output weights, as shown in the last row at the bottom of Figure 9 in the main text), giving rise to a sum squared error of 2.5 between the correct sparrowhawk output and the average bird pattern. This happens because the input features of the sparrowhawk project fully onto dimensions 1 and 3 of the knowledge already in the network, and these dimensions together capture the average properties of the existing birds.

From this starting point, learning proceeds in each of the different interleaving conditions much as in the one-hot input setup, but with adjustments to the dynamics due to the auto-associative setup, in line with those we observed in Figure 6b in the main text in considering how the auto-associator learned the original eight-item data set from scratch. As shown in the top row of Figure 4, focused presentation of only the sparrowhawk leads to rapid learning of this item, at the cost of extensive interference with the already known birds, lesser interference with the fish, and no interference with the trees or flowers, as in the feed forward network case. Furthermore, and also as before, fully interleaved learning results in reduced interference with existing items, at the cost of a slowdown in fully capturing the properties of the sparrowhawk until the new dimension that separates the sparrowhawk from the existing birds is learned. Also as before, the time to learn this new dimension is exactly what it would have been had the network be learning the whole nine-item data set from scratch. In addition, as Figure 5 shows, similarity weighted interleaving using the same presentation rates per epoch as in the feed forward network produces virtually identical results compared with full interleaving, with the same reduction in the total amount of interleaved learning required. Finally, in the control condition the relative slowdown in the time to learn the new dimension is still strictly a matter of the density of exposure to the relevant items (the sparrowhawk and the other two birds) times the number normalized training epochs.

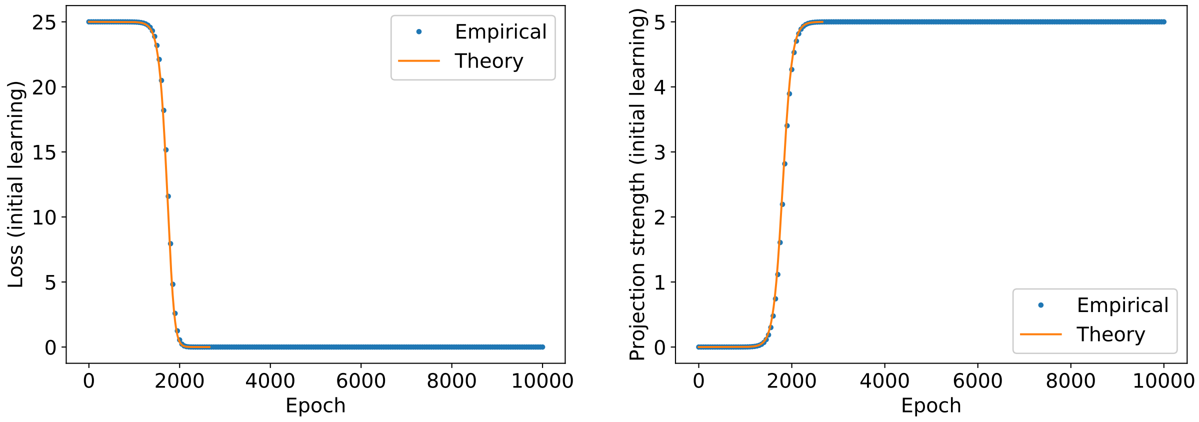


(a) Loss

(b) Mode alignment

*Figure 1*. Match between theory (Eq. 1) and empirical initial learning of a single mode. (a) shows the loss (squared error) of the network's outputs, and (b) shows the alignment (i.e. the quantity $ab$).

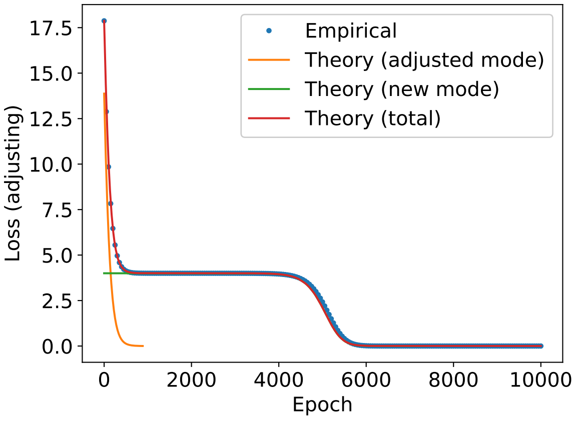


(a) Loss

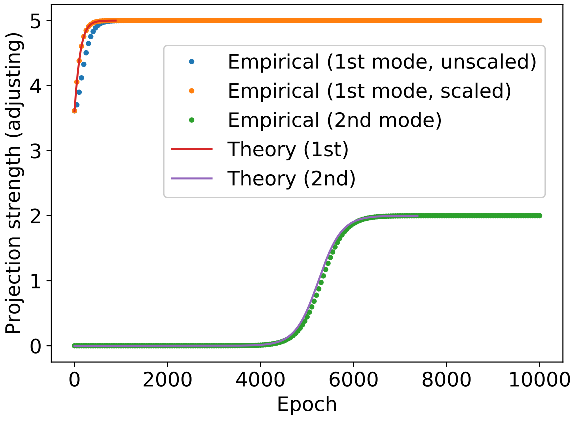


(b) Mode alignments

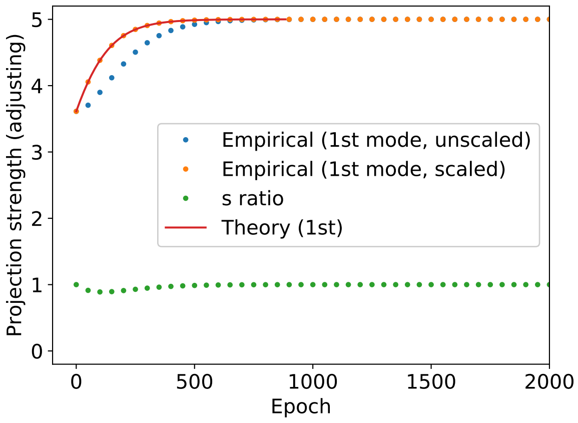


(c) Adjusting mode alignment detail

*Figure 2*. Match between theory (Eq. 1) and empirical adjustment of one mode while simultaneously learning an orthogonal new mode. (a) shows the theoretical loss due to each component as well as the total, showing an extremely close match between theoretical and empirical total loss. (b) shows the alignments of the modes, showing a slight discrepancy in the alignment of the first mode, which is due to a transient decrease in the singular value. (c) shows this discrepancy in the alignment in more detail, including the ratio of the empirically observed $\hat{s}$ to $s$, and that when the alignment is scaled by the inverse of this ratio it matches the theory exactly. (Singular values: $s_{old} = 5$, $s_{new} = 2$.)

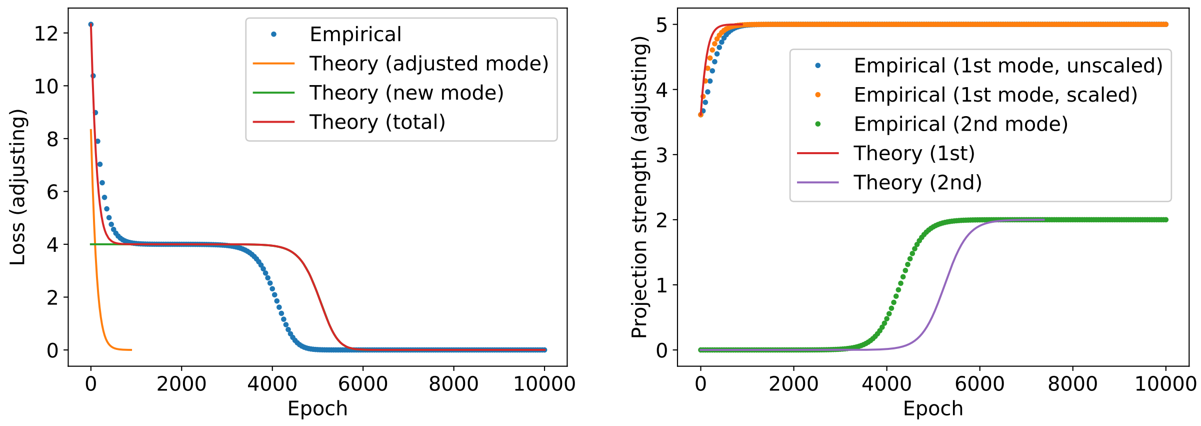


(a) Loss

(b) Mode alignments

*Figure 3*. Match between theory (Eq. 1) and empirical adjustment of one mode while simultaneously learning a new mode which is partially aligned with the old mode. The theory curves in the partially aligned case are the same as in the orthogonal case, showing the slight slow down in adjusting the old mode and speed up in learning the new mode.(Singular values: $s_{old} = 5$, $s_{new} = 2$.)

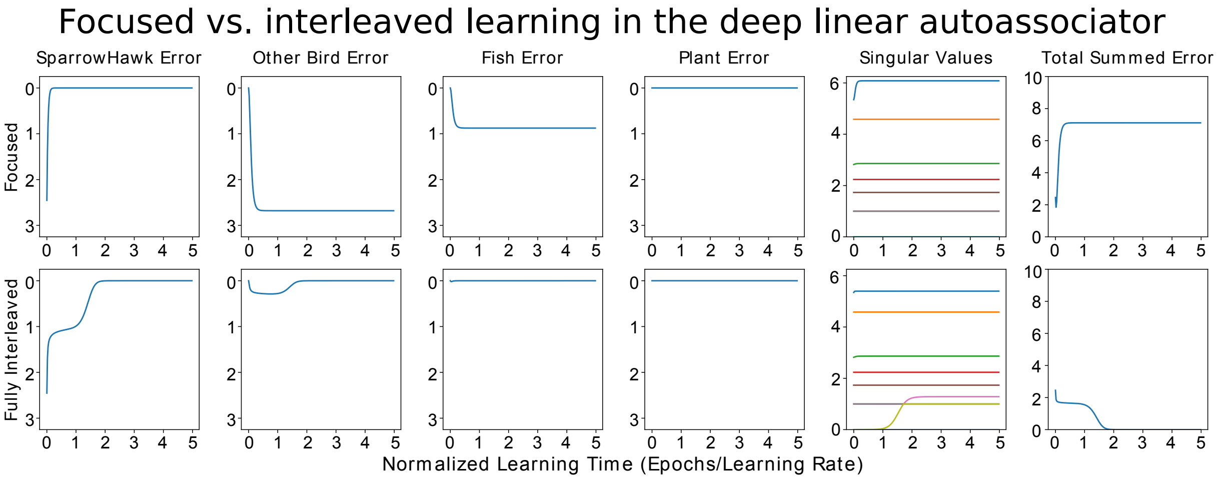


*Figure 4*. Focused vs. Interleaved Learning of a new item in the deep linear auto-associator. Conventions as in Figure 8 of main text. Note that, initially, the sum squared error on the SparrowHawk is only 2.5. During focused learning, this error is quickly eliminated, at the cost of The network is activating the average known bird, exactly as after the first stage of learning with frozen output weights in the one-hot input case. As before, dashed line shows expected learning time for the added dimension when all items are learned in a fully interleaved fashion without any prior learning. Thus, learning the new dimension still requires the same amount of interleaved learning of the birds as it would have required to learn the dimension without any prior learning.

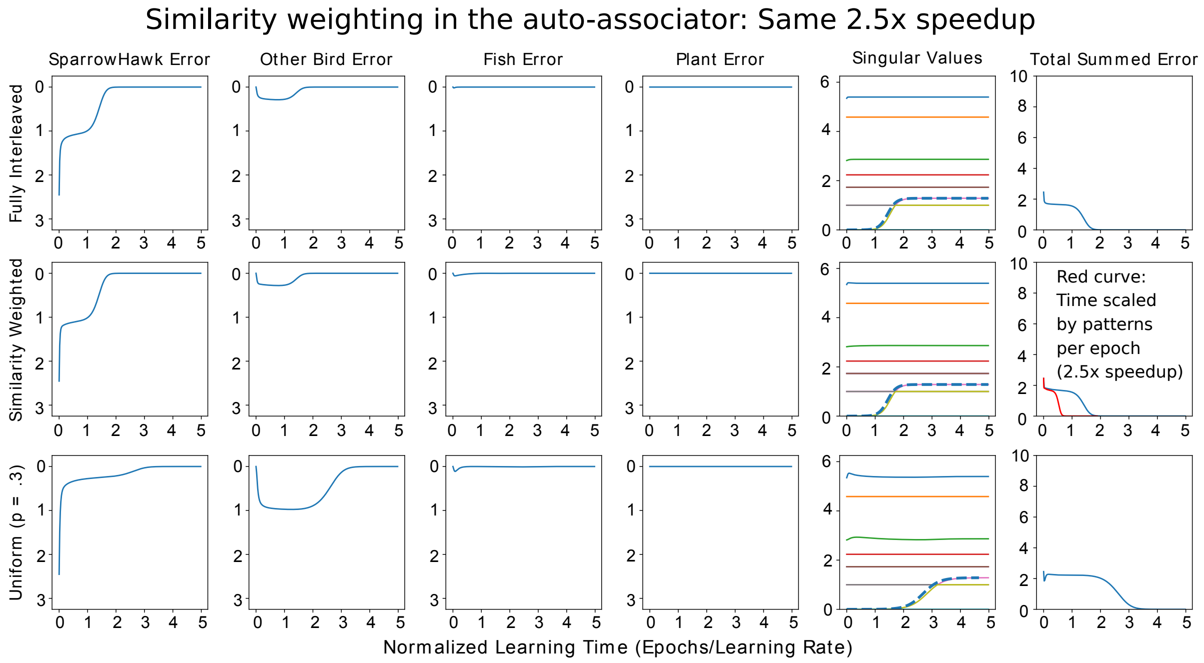


*Figure 5*. Similarity weighted Interleaved learning in the linear auto-associative network (middle) compared with Full Interleaving (top) and the a control condition in which each known item is presented .3 times per epoch (bottom). Conventions as in Figure 10 of main text. Average number of pattern presentations per epoch is 9 for Full interleaving, and 3.4 for the other two cases, or 38% of the pattern presentations with full interleaving. Dashed lines shows expected learning time for the new dimension when all items are learned in a fully interleaved fashion without any prior learning (top two rows), or when scaled for the average number of presentations of relevant items in the control condition (bottom row).
